# A high fermentable fiber Western diet reduces indole levels

**DOI:** 10.64898/2026.01.27.702025

**Authors:** Medha Priyadarshini, Julianne Amy Jorgensen, Sophia R. Chang Stauffer, Lina Issa, Nupur Pandya, Chioma Nnyamah, Kai Xu, James E Boyett, Parneet Kular, Aditi Mhatre, Vishal H Brahmbhatt, Jack A. Gilbert, Md Wasim Khan, Barton Wicksteed, Yang Dai, Brian T Layden

**Affiliations:** Department of Medicine, Division of Endocrinology, Diabetes, and Metabolism, University of Illinois Chicago (UIC), Chicago, IL, U.S.A; Department of Pediatrics, University of California San Diego (UCSD) School of Medicine, La Jolla, California, U.S.A; Scripps Institution of Oceanography, UCSD, La Jolla, California, U.S.A; Department of Oncology, Barbara Ann Karmanos Cancer Institute, Wayne State University, Detroit, MI, U.S.A; Richard and Loan Hill Department of Biomedical Engineering, College of Engineering and Medicine, UIC, Chicago, IL, U.S.A; Jesse Brown Veterans Affair Medical Center, Chicago, IL, U.S.A

**Keywords:** gut microbiome, dietary fiber, indole, indoxyl sulfate, insulin resistance, western diet

## Abstract

Changes in gut microbiota composition due to diet impact health. Fiber-rich diets promote beneficial microbiota and reduce the risk of metabolic diseases, while low-fiber, calorie-dense diets are linked to dysbiosis and increased disease risk. This study examines the effects of a Western diet (WD) and explores dietary fiber supplements as potential modifiers of those effects. 10-week-old C57Bl/6J male mice were fed control (low-fat) or WD (high-fat, high-sucrose) containing 0% fermentable fiber (FF) or WD supplemented with 20% FF (fructooligosaccharides, FOS; guar gum, GG, or pectin, Pec). After 19 weeks, analysis of the cecal metagenome using whole-genome shotgun sequencing, metabolome by untargeted and targeted LC-MS/MS, and tissue RNA and protein expression by RT-PCR and immunoblotting was undertaken. WD-FF reduced metabolic derangements from WD while also improving GM diversity and altering cecal metabolites, particularly tryptophan metabolism. A profound increase in cecal indole levels (targeted metabolomics) was noted in WD vs WD-FF groups. As the primary indole-oxidizing enzyme, CYP2E1 generates indoxyl sulfate, which contributes to oxidative stress and a leaky gut. Mice on WD displayed higher expression of *Cyp2e1* mRNA in the gut. In the liver, the levels of both CYP2E1 protein and mRNA were higher in the WD group compared to the WD-FOS group, with protein levels also higher than in the WD-Pec group and mRNA levels higher than in the WD-GG group. mRNA expression of markers of oxidative stress, inflammation, and leaky barrier was significantly higher in the liver and intestine of the WD vs the WD-FF groups. FFs reduced high plasma indoxyl sulfate levels (except in WD-GG), and boosted short-chain fatty acids and indole acetic acid. Our data suggest that WD disrupts GM tryptophan metabolism, possibly by altering the balance between indole-producing and utilizing gut bacteria. Dietary fiber supplementation exerts protective effects, in part, by mitigating this imbalance.

## Introduction

Non-communicable diseases (NCDs) such as overweight and obesity, diabetes, cardiovascular diseases, and chronic kidney diseases have become leading causes of death in high-income countries ^1^. Research in animal models and human cohorts has linked many of these NCDs to the gut microbiome (GM) ^2^. While the environment and genetics influence the GM, diet has emerged as a key factor in determining the composition and function of GM ^3^ with westernization of the diet linked to NCD in mice ^4^ and in humans ^5, 6^. A Western-style diet (WD) produces shifts in gut ecology, reducing diversity, depleting beneficial commensal taxa, and promoting the growth of facultative anaerobes ^7, 8^. Due to these shifts, microbial metabolites are either depleted (such as short-chain fatty acids, SCFAs^9^) or produced at harmful levels (such as indole, a precursor to the uremic toxin indoxyl sulfate^10^), which can lead to adverse health effects^11^. Clinical investigations aim to restore gut microbial homeostasis and/or replenish depleted microbial metabolites through fecal microbial transplants from healthy donors to obese individuals^12, 13^or, more simply, through modified diets that cultivate a diverse and functional GM^1^.

In this context, dietary fibers, nondigestible plant carbohydrates, can shape the structure and function of GM. Notably, WD has a low fiber content that falls short of the recommended daily dietary fiber intake of 25–35 g in high-income countries ^14^. This lack of dietary fiber has been postulated as a key contributor to NCDs ^15^. In fact, several epidemiological and interventional studies have shown that high dietary fiber intake is associated with reduced risk of all-cause mortality and NCDs ^1, 16^, reflecting the metabolic benefits conferred by dietary fiber.

Based on their fermentability, dietary fibers are classified into two groups: insoluble forms, which primarily affect fecal mass and intestinal motility, and soluble forms, which are metabolized by gut bacteria ^17^. Soluble dietary fiber is fermented by gut bacteria, with SCFAs as a key byproduct ^18^. Dietary fibers can promote microbial diversity through niche specialization and cross-feeding of byproducts with non-fermentative species ^19^. These diverse gut bacteria participate in multiple functions, including the synthesis of metabolites (secondary bile acids, vitamins ^18^), the protection of the mucus layer ^20, 21^, and the maturation and priming of the immune system ^22, 23^. Mainly through these properties and the promotion of specific metabolites relevant to host physiology, the efficacy of fermentable fibers, such as inulin, oligofructose, guar gum, and pectin, has been explored in rodent disease models and human cohorts ^1^. However, it is challenging to draw a comparison in fiber efficacies from rodent studies, mainly due to differences in models (such as the type and duration of high-fat or fiber diet feeding). Furthermore, most studies have focused on fiber-induced alterations in specific metabolites, such as SCFAs and bile acids. For human studies, these data are often confounded by interindividual variability in gut microbial responses to different types of fibers.

Thus, in the current highly controlled studies, we have explored and compared the physiological effects of 3 different purified fermentable fibers, specifically, fructo-oligosaccharide (FOS), pectin (Pec), and guar gum (GG), in a western diet (WD) murine model. We observed that fiber supplementation provided a predictable and remarkable recovery from WD-induced metabolic derangement, with the 3 fibers differing in the robustness of their beneficial effects. Diet-dependent shifts in specific gut bacterial strains were observed, with an overall preference for proteolytic fermentation in WD-associated gut bacteria compared to saccharolytic fermentation in fiber-supplemented groups. Markedly increased cecal indole levels, and associated increase in plasma indoxyl sulfate level in the WD-fed group, contributed to gut and liver inflammation. Dietary fiber supplementation restricted the increase in these metabolites, mitigating WD-induced metabolic derangement.

## Materials and Methods

### Mouse Husbandry

Animal care and study protocols were approved by the University of Illinois Chicago Animal Care and Use Committee and performed in accordance with the Guide for the Care and Use of Laboratory Animals. Male C57BL/6J (age 9 weeks) were procured from the Jackson Laboratory and group-housed (5/cage) on a 12-hr light:dark cycle in a temperature- and humidity-controlled, specific pathogen-free barrier facility with ad lib access to autoclaved food (Inotiv 7912) and water. After 1-week acclimation period, mice were randomly assigned to specially formulated diets-control (10% Kcal fat; 17% protein; 73% carbohydrate, D19010908, Research Diets), western (40% Kcal fat; 17% protein; 43% carbohydrate, D19070209, Research Diets) or western diet supplemented with 20% fiber content by weight as FOS (Nutraflora FOS powder, NOW foods, Bloomingdale, IL, D19070210, Research Diets), pectin (TIC Gums, D19070211, Research Diets), guar gum (TIC Gums, D19070212, Research Diets). For all diets, standard corn starch was replaced by non-fermentable amioca starch ^24^. The fiber diets were macronutrient-matched to the WD, with similar but not identical kilocalories in some instances (Supplementary Table S1). All the diets had the same amount of insoluble fiber, cellulose. At the end of the study period, cecal material, blood, liver, intestine, and adipose tissue were collected, flash-frozen in liquid nitrogen, and stored at -80 °C until processing.

### Metabolic Assays and Measurements

Body weight and ad lib glucose were monitored weekly. Mice were fasted for 16 hr and blood (from the tail vein) was collected at regular time points during the study for measurement of glucose (OneTouch Ultramini glucometer, Lifescan, Malvern, PA), insulin (ELISA, ALPCO, Salem, NH), triglycerides, and non-esterified free fatty acids (NEFAs) (Wako Diagnostics, Richmond, VA). Mice were fasted for 16 hours and 6 hours prior to receiving an intraperitoneal injection of glucose (2 g/kg) or insulin (1.0 U/kg), respectively, for glucose and insulin tolerance tests. The tests were conducted as described previously ^25^.

#### Body composition and metabolic rate

Whole-body lean, fat, and free fluid mass were quantified via NMR using a Minispec LF50 Body Composition Analyzer (Bruker, Billerica, MA) at 8 and 16 weeks following dietary intervention. Energy expenditure, VO_2_, VCO_2_, respiratory exchange ratio, food intake, and activity were measured using the Mouse Promethion Continuous caging system (Sable Systems International, Las Vegas, NV). Data are presented as hourly averages for each metabolic parameter and analyzed using ANCOVA performed with the CalR tool (https://calrapp.org) ^26^.

#### Analyte measurements

Plasma liver triglyceride by reagents from Wako Diagnostics ^27^. Plasma and cecal samples at the terminal time point were analyzed via LC-MS/MS using both an untargeted approach and a targeted approach for SCFAs and tryptophan metabolites, as described previously ^25^. For the latter, plasma and feces were treated with methanol. L-Kynurenine sulfate-13C10 (Kyn-13C10), tryptophan-d5 (Trp-d5), serotonin- d4 (5-HT-d4), tryptamine-d4 (Try-d4), indole-13C4 (I-13C4) and indoxyl-3a,4,5,6,7,7a-13C6 sulfate were used as internal standards (IS). LC-MS/MS measurements were performed at the Mass Spectrometry Core facility of the UIC Research Resource Center.

### RNA extraction and quantitative real-time PCR

1 μg purified extracted RNA (RNeasy Mini Kit, Qiagen) was reverse transcribed using qScript Reverse Transcriptase (Quanta Biosciences) and amplified using iTaq Universal Syber Green supermix (BioRad) with primers listed in Supplementary Table S2. The expression of individual genes was normalized to the housekeeping gene β-actin or Gapdh and presented as a fold change using the 2^ΔΔCt^ algorithm.

### Immunoblotting

Protein lysates were prepared using RIPA buffer (Cell Signaling) supplemented with protease inhibitor cocktail and phosphatase inhibitor cocktail (Thermo Fisher Scientific). Equal amounts of total protein (estimated using Bradford reagent, Bio-Rad) for each sample were separated on sodium dodecyl sulfate polyacrylamide gel electrophoresis (15%) and transferred to polyvinylidene difluoride membranes. The membranes were probed with the use of CYP2E1 (Proteintech #19937-1-AP, 1:5000), SULT1A1 (Proteintech #10911-2-AP, 1:1000), HO-1 (Proteintech #10701-1-AP, 1:3000), NQO-1

(CST #D6H3A, 1:1000), GAPDH (CST #14C10, 1:1000). Proteins were visualized using the Amersham ECL western blotting detection kit and an image analyzer (Bio-Rad).

### Histology and histological scoring

Formalin-fixed and paraffin-embedded tissues were sectioned at 5 μm by the Research Histology and Tissue Imaging Core of the University of Illinois Chicago and stained with Hematoxylin & Eosin (Vector labs) and imaged using Olympus BX51 light microscope. The sizes of the adipocytes were analyzed using the ImageJ version 1.53a software (https://imagej.nih.gov/ij) with the Adiposoft version 1.16 plug-in (https://imagej.net/adiposoft) to calculate adipocyte size and number of adipocytes per image ^28^.

### Shotgun metagenomic sequencing and data analyses

Total genomic DNA was extracted using a standardized protocol and quantified prior to library preparation. Libraries were constructed using the KAPA Hyper Plus kit and sequenced on the Illumina NovaSeq 6000 platform. Raw sequence files were demultiplexed using BaseSpace (Illumina, CA, USA); then adapter-trimmed and quality-filtered to remove low-quality reads and bases, and host-filtered for quality control ^29^. Host-depleted, quality-filtered reads were uploaded to Qiita for per-sample metagenomic microbial classification using the Web of Life (WoL) toolkit (Woltka). Direct genome alignment against the WoL reference database ^30, 31^ was performed using Bowtie2 as the backend ^32^. Reads mapping to microbial reference genomes were counted to generate a feature table of samples by microbial genome identifiers, referred to as operational genomic units (OGUs). Reads mapping to multiple genomes were fractionally assigned (1/k per genome), and genome counts were summed and rounded to generate a final count matrix, saved as a BIOM file and a feature table for downstream analyses, and processed in R ^33, 34^.

Alpha diversity was calculated from the taxonomic feature table using richness- and evenness-based metrics, including the Shannon diversity index. Prior to diversity analyses, samples were normalized to account for differences in sequencing depth using rarefaction. Beta diversity was assessed using multiple distance metrics, including Bray– Curtis dissimilarity, as well as weighted and unweighted Unifrac distances ^35, 36^. Differential analysis of microbial taxa: Gene IDs from BIOM files were mapped to genome IDs and taxa using lineages from Web of Life (https://biocore.github.io/wol/). Taxonomic summaries were generated at the phylum to species levels. Heatmaps were constructed using the ComplexHeatmap R package (https://bioconductor.org/packages/release/bioc/html/ComplexHeatmap.html).

Differential analysis of functional genes: Gene IDs were mapped to UniRef protein IDs from the UniProt database and then to KEGG ortholog (KO) IDs. Higher level summaries of the predicted orthologous functions were created using KEGG pathway, module and BRITE hierarchical annotations ^37, 38^.

Enrichment analysis of functional gene orthologs: The enrichment or overrepresentation of differentially abundant functional gene orthologs in the various functional gene groups, i.e. pathways, modules and BRITE categories, was determined using Fisher’s Exact test in R. Briefly, lists of differentially abundant functional gene orthologs were obtained from the tests of the various pairwise comparisons based on a q value, i.e. false discovery rate (FDR) corrected p value, less than 0.05. For each model term or pairwise comparison to be considered in the enrichment analysis, a minimum of 10 significantly different orthologs was required. The enrichment of these significant orthologs for each model term or pairwise comparison, relative to all detected orthologs, was then evaluated across all identified KEGG pathways, modules, and BRITE categories at levels 1, 2, and 3.

### Untargeted metabolomics

Basic processing: Feature detection and basic quantitation of raw LC-MS data were performed using the OpenMS toolkit ^39^. Briefly, raw LC-MS data were filtered to remove spectra before/at solvent peak and after wash phase. The features were then detected using peak shape parameters optimized for the separation conditions and an isotopic filtering model consistent with non-halogenated metabolites. Features were quantified using the integrated area of extracted ion chromatograms.

The resulting feature file for each sample was then aligned and combined into a single consensus feature file for each acquisition mode, i.e., positive and negative. The intensity of the features in consensus feature files was then normalized to internal standards added to the samples prior to data acquisition.

Differential analysis: Prior to differential analysis, the consensus feature table was filtered to remove features that were present in less than 50% of the samples. The values were log 10 scaled, z-scored, and normalized using quantile normalization.

### Statistical analyses

Statistical differences in microbial community composition were evaluated using permutational multivariate analysis of variance (PERMANOVA) ^40^. Univariate comparisons were conducted using two-tailed t-tests when normality assumptions were met or else Wilcoxon Signed Rank tests, with multiple testing controlled using the Benjamini–Hochberg false discovery rate; adjusted p values < 0.05 were considered statistically significant.

Differential analyses of orthologous gene features were performed using the software package edgeR on raw sequence counts in a similar fashion as taxonomic summaries. ^41, 42^. Prior to analysis, the summary tables were filtered to remove any groups that accounted for less than 1000 of the total sequence counts or were absent in more than 30% of the samples. The orthologue data were filtered to remove orthologues that had less than 100 total counts or were absent in more than 30% of the samples. Data were normalized as counts per million. Pairwise comparisons between diets were performed using the “exactTest” function in edgeR.

For metabolomics data, differential gene expression analysis was performed using the limma package ^43^. Q values were calculated using the Benjamini-Hochberg FDR correction ^44^. Significant orthologs or groups were determined based on an FDR threshold of 5% (0.05).

All other data are expressed as means±SEM and analyzed by Student’s two-tailed unpaired *t* tests, one-way ANOVA or two-way ANOVA with post hoc tests (GraphPad Software 10.0) as applicable. *P*<.05 was considered significant.

## Results

### Fermentable Fiber Alleviates WD-Induced Metabolic Syndrome

In this study, we used a 19-week exposure to three different purified fermentable fibers supplemented in the obesogenic WD to assess their effects on WD-induced metabolic derangement (Figure 1A). Mice exposed only to the WD (deprived of any fermentable fiber, Suppl. Table 1) served as the control. Relative to the WD, WD supplemented with 20% (w/w) fermentable fiber (FF) caused a marked reduction in weight gain (Figure 1B-C). Among the three FFs, the guar gum-supplemented diet (WD-GG) resulted in the greatest decline in weight gain, while the least decline was observed with fructooligosaccharide (WD-FOS) supplementation (Figure 1C). Without affecting the lean mass, all FFs restricted gain in fat mass (Figure 1D), diminishing WD-induced adiposity as assessed by the subcutaneous and epididymal fat pad weights (Figure 1F) and significantly reduced adipocyte size (Figure 1G). The groups were also characterized by indirect calorimetry for food intake, energy expenditure, VO_2_, VCO_2_, and respiratory exchange ratio, and the data were analyzed by ANCOVA. No significant differences were noted in the food intake (Figure 1E). After adjusting for fat mass and total mass, energy expenditure and VO_2_ exhibited near-significant trends in the dark period (∼p=0.06, Suppl. Figure 1), while VCO_2_ reached significance in the dark period (p=0.039 with fat mass and p=0.053 with total mass as covariate, Suppl. Figure 1). This suggested a diet-induced metabolic shift, independent of adiposity, with a higher preference for fat as fuel in WD-fed mice (reflected by higher VO_2_). FFs also improved glycemic control. Ad lib (Figure 1H) and fasting blood glucose (Figure 1I), HOMA-IR (Figure 1K), and glucose intolerance (Figure 1L) were significantly lower in FF supplemented groups. In one exception, the 8-week fasting glucose level was higher in the WD-GG group (Figure 1I). In the absence of any other metabolic anomaly (unchanged fasting insulin level (Figure 1I), TGs and NEFAs (Figure 1J), and fasting glucose at 19 weeks (Figure 1L)), this could be due to a temporary stress response in WD-GG mice. Compared with WD-fed mice, WD-Pec mice displayed higher fasting TGs and NEFAs at 8 weeks into the dietary intervention, and TGs remained significantly higher under ad lib conditions even at 19 weeks (Figure 1J). Elevated circulating TGs and NEFAs may reflect increased VLDL secretion and adipose lipolysis. However, a normal insulin sensitivity and glucose tolerance indicate preserved metabolic control. FFs enhanced insulin sensitivity (Figure 1M). As compared to WD-fed mice, fiber-supplemented groups required less insulin to maintain glucose levels during GTT (Figure 1L). Confirming improved insulin action, the rate of glucose disappearance (Kitt) was significantly higher in WD-FOS and WD-GG groups and trended higher in WD-Pec group (p=0.08) (Figure 1M).

**Figure 1.**
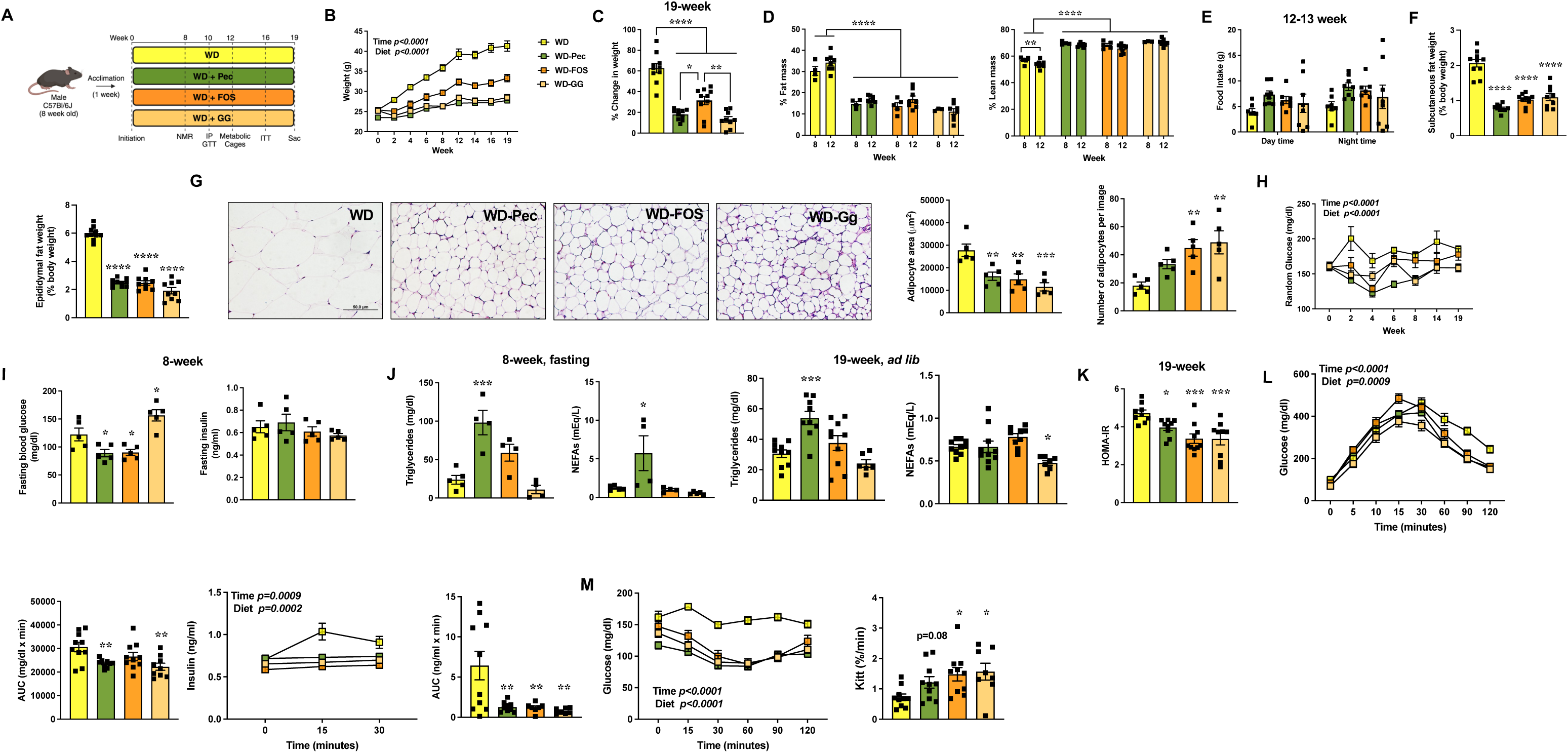
Dietary fiber supplementation alleviates the metabolic derangements associated with Western diet-induced obesity. A) Schematic of the experimental design. B) Body weight. C) Percent change in body weight at 19 weeks with respect to baseline. D) Fat mass and lean mass at the indicated times. E) Food intake at the 12-13 week time point. F) Weights of subcutaneous and epididymal adipose tissues. G) Representative images of H & E-stained subcutaneous adipose tissue, average area of individual adipocytes, and number of adipocytes per image. H) Ad lib glucose levels measured throughout the study. I) 8-week fasting glucose and insulin levels. J) 8-week fasting and 19-week ad lib, triglycerides and NEFA levels. K) 19-week HOMA-IR. L) IP-GTT measured after 10 weeks on the diet, with corresponding AUC and insulin levels, and AUC during the first 30 minutes of the test. M) ITT with glucose clearance. Data are mean ± SEM; n=7-10 per group for B, C, F, H, K, L, M; n=3-5 per group for week 8 and n=8 per group for week 12 measurements for D; n=7-8 per group for E; n=4-5 per group for G; n=4-5 per group for week 8 and n=6-10 per group for week 19 measurements for J. Data were analyzed by 2-way ANOVA Mixed-effects analysis with Bonferroni post-hoc tests for B and E, Dunnett for D, L and M or 1-way ANOVA with Tukey’s post-hoc analysis for C and F, Dunnett for G and I-K, and insulin and glucose AUCs presented in panel L with pair-wise comparisons (WD vs each WD-FF group). NEFA, non-esterified fatty acid; HOMA-IR, homeostasis model assessment of insulin resistance; IP-GTT, intraperitoneal glucose tolerance test; AUC, area under the curve; ITT, insulin tolerance test; ANOVA, analysis of variance. *P < 0.05; **P < 0.005; ***P < 0.001; ****P < 0.0001.

Additionally, to provide a physiological baseline, a distinct group of mice fed a low-fat control diet (CD) was compared to the WD-fed group (Suppl. Figure 2A). As expected, CD reduced weight gain and adiposity (Suppl. Figure 2B-E), supported better glycemic control (Suppl. Figure 2F, G & J), and insulin sensitivity (Suppl. Figure 2G & J). Fasting and ad lib TGs and NEFAs did not differ significantly in CD and WD-fed groups (Suppl. Figure 2H & I).

### Fermentable Fibers Alter Cecal Microbiota Regulating Tryptophan Metabolism

Shotgun metagenomic sequencing of the cecal contents revealed a strong influence of FF supplementation on bacterial diversity and composition. Alpha diversity (Shannon entropy) increased significantly in FF groups compared to the WD-fed group (Suppl. Figure 3A). In contrast, CD and WD, with no FF, showed similar alpha diversity (Suppl. Figure 3A). To determine shifts in community structure (beta diversity), Bray-Curtis dissimilarity index, Weighted Unifrac, and Unweighted Unifrac (Figure 2A) were used. The Bray-Curtis dissimilarity index (which considers the relative abundance of species), as well as Weighted and Unweighted Unifrac distance (which consider phylogenetic distances between samples, weighting relative abundance or presence and absence of species, respectively), analyses revealed significant separation between the WD and WD-FF fed groups (PERMANOVA, FDR corrected p-value < 0.005). CD and WD-fed groups also exhibited significant separation in all the indices of beta diversity (PERMANOVA, FDR corrected p-value < 0.01) (Suppl. Figure 3B-D). Overall, these data indicate diet-driven shifts in gut microbial community composition, reflected in both the number and abundance of unique and common species.

**Figure 2.**
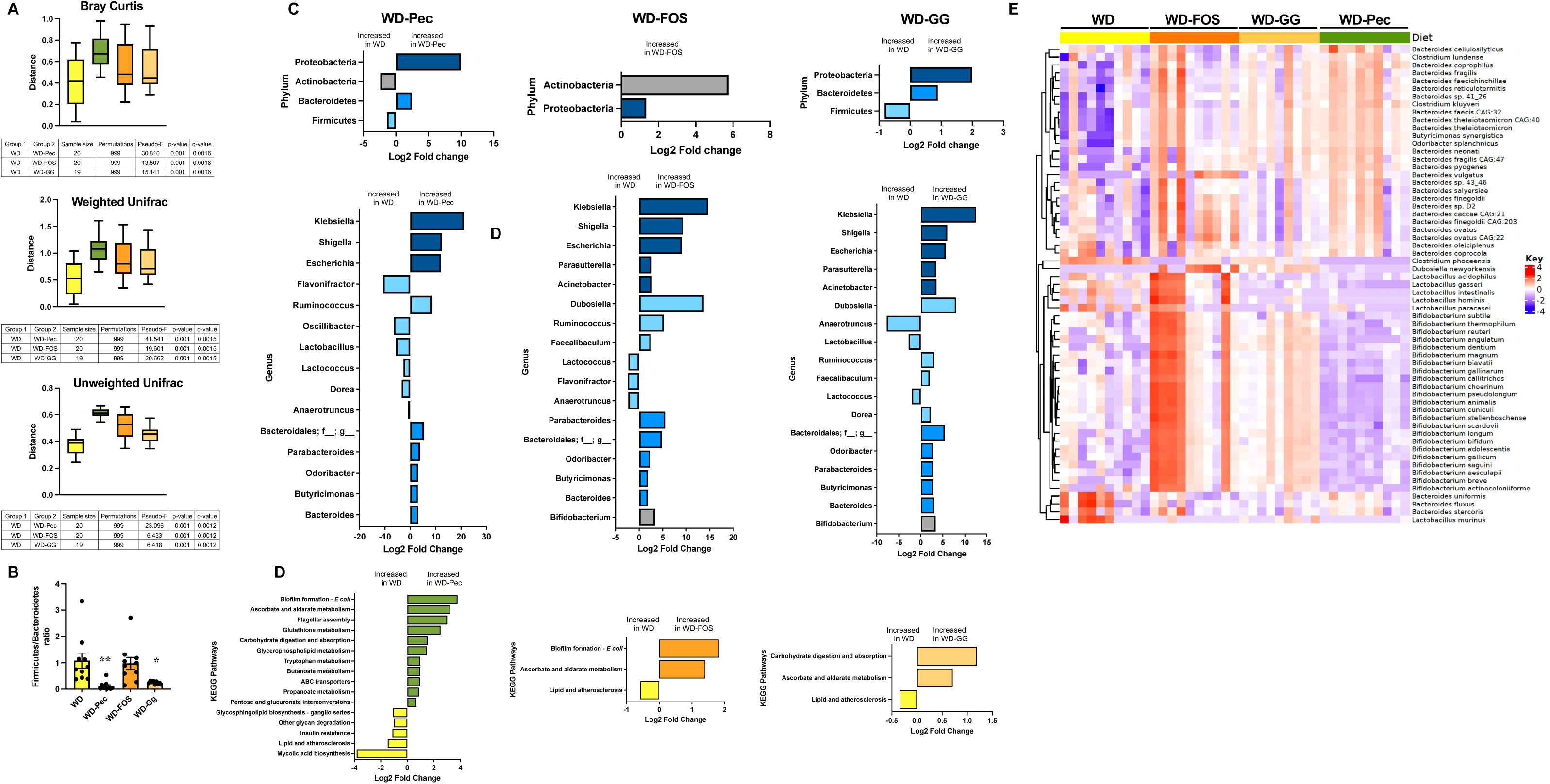
Dietary fiber supplementation alters the cecal gut microbiome associated with the Western diet. A) Beta diversity: Bray-Curtis, Weighted, and Unweighted Unifrac distances. B) Firmicutes/Bacteroidetes percentage ratio presented as mean ± SEM. C) Bar plots at two taxonomic levels (phylum and genus) depicting log2 fold changes for pair-wise comparisons between samples. D) Bar plots depicting log2 fold changes for relevant KEGG pathways that were significantly enriched in the treatment groups and were also significant in the differential analyses. E) Heatmap of species with significantly abundant changes from genus *Bifidobacterium*, *Bacteroides, Lactobacillus, Butyricimonas, Clostridiales, Odoribacter, Dubosiella*. Heatmap was constructed using the ComplexHeatmap R package using log2-scaled and z-scored normalized abundances. Each row represents a species, and each column represents a sample. n=9-10 per group. Data analyzed by PERMANOVA for A, and 1-way analysis of variance with Bonferroni post-hoc tests for B. For C and D, pairwise comparisons between diets were performed using the “exactTest” function in edgeR. Adjusted P values (Q values) were calculated using the Benjamini-Hochberg false discovery rate correction and considered significant with a Q value <0.05.

Obesogenic WD and FFs led to unique phylum- and genus-level differences. In our study, the Firmicutes/Bacteroidetes (F/B) ratio decreased significantly in the WD-Pec and WD-GG groups (Figure 2B). No difference in this ratio was seen between WD and WD-FOS fed mice (Figure 2B). The F/B ratio is frequently used as an indicator of obesity-associated dysbiosis, with *Firmicutes* being selected by a high-fat diet and *Bacteroides* by fiber ^45^. In our data, FFs, Pec, and GG promoted Bacteroidetes expansion, whereas FOS selected *Actinobacteria*. These fiber-specific selections could have contributed to the changes observed in the F/B ratio. Surprisingly, the F/B ratio was significantly higher in CD-fed mice compared to the WD group (Suppl. Figure 3E). CD consumption led to the expansion of *Firmicutes*. The fermentable fiber-deprived CD used in the study serves as a low-calorie control for the WD, but it contains the same amount of cellulose as the WD.

Expansion of *Firmicutes* in CD-fed mice may contribute to efficient energy extraction and the degradation of insoluble fiber ^46^. Furthermore, our data indicate that the F/B ratio does not necessarily correlate with body mass or metabolic disease, a finding also reported by others, suggesting that this ratio lacks the nuance to capture the complexity of the gut microbiota ^45, 47^. Among other phyla, all three FFs in the presence of WD stress led to a bloom in *Proteobacteria*, as reflected by an increase in several genera, including *Klebsiella*, *Shigella,* and *Escherichia,* which belong to this phylum and the family *Enterobacteriaceae* (Figure 2C). WD also increased the proportion of *Anaerotruncus*, *Flavonifractor*, and *Lactobacillus* (phylum *Firmicutes*), which have been shown to be associated with obesity and consumption of high saturated fats (^48^). In accordance with earlier reports, fiber degraders, such as *Butyricimonas*, *Odoribacter*, and *Parabacteroides*, were increased by all the FFs, while *Bifidobacterium* expanded specifically in the presence of FOS and guar gum (Figure 2E) ^49–51^. The genome data were consistent with enrichment of pathways related to lipid metabolism (lipid and atherosclerosis, insulin resistance, etc.) in the WD groups and with enrichment of pathways involved in biofilm formation in the WD-FF groups (Figure 2D). Other pathways enriched in WD-FF groups indicated changes in oxidative stress (ascorbate and aldarate metabolism), SCFA production (butanoate and propanoate metabolism), and tryptophan metabolism. Thus, focusing on a subset of bacterial species contributing to these pathways (Figure 2E), we observed the expected specific pattern of complex carbohydrate degraders and SCFA producers, such as *Bacteroides cellulosilyticus*, *Bacteroides vulgatus*, *Bacteroides ovatus*, *Bacteroides finegoldii*, and *Odoribacter splanchnicus,* which increased in proportion in all FF groups with a higher prevalence in WD-Pec group ^52^. *Bacteroides* and *Lactobacillus* species associated with gut inflammation increased uniquely in WD mice ^53–55^. FOS was specifically shown to significantly increase the proportion of several *Bifidobacterium* species. These species-level differences were less defined between CD and WD-fed mice, except for the greater proportion of *Bifidobacterium* species in the CD group (Suppl. Figure 3H). Reduced levels of free fatty acids and simple sugars in the intestine of CD-fed mice would likely prevent the inhibition of the growth of *Bifidobacterium* sensitive to these metabolites ^56, 57^.

Importantly, most of the species that increased under FF fiber consumption have been associated with shifts in bacterial tryptophan metabolism ^58–61^. To determine if these microbial shifts are reflected in the plasma metabolome of experimental mice, we conducted untargeted metabolomics, which detected a total of 4085 metabolites in the plasma from CD, WD, and WD-FF-fed mice. As shown in the PCA, the groups did not cluster based on diet, except for the WD-GG group, which exhibited the greatest separation from the other groups (Figure 3A and Suppl. Figure 4A). Pairwise comparisons between diet groups identified 1 feature in the WD-Pec/WD group, 2 in the WD-FOS/WD group, 9 in the WD-GG/WD group (Figure 3B), and 2 in the WD/CD group (Suppl. Figure 4B) that showed significant differences between the two groups (FDR corrected p-value <0.05, and log2fold change >0). Most of these features could be annotated to more than one metabolite, which were related to amino acid and lipid metabolic pathways, including dipeptides (Prolyl-threonine and γ-glutamyltyrosine), tryptophan metabolism (kynurenic acid, quinaldic acid), decanoylcarnitine, and lyso-PE. Among these, tryptophan metabolism emerged as a commonality among the bacterial species selected by WD-FF diets, and plasma metabolites highlighted by untargeted metabolomics.

**Figure 3.**
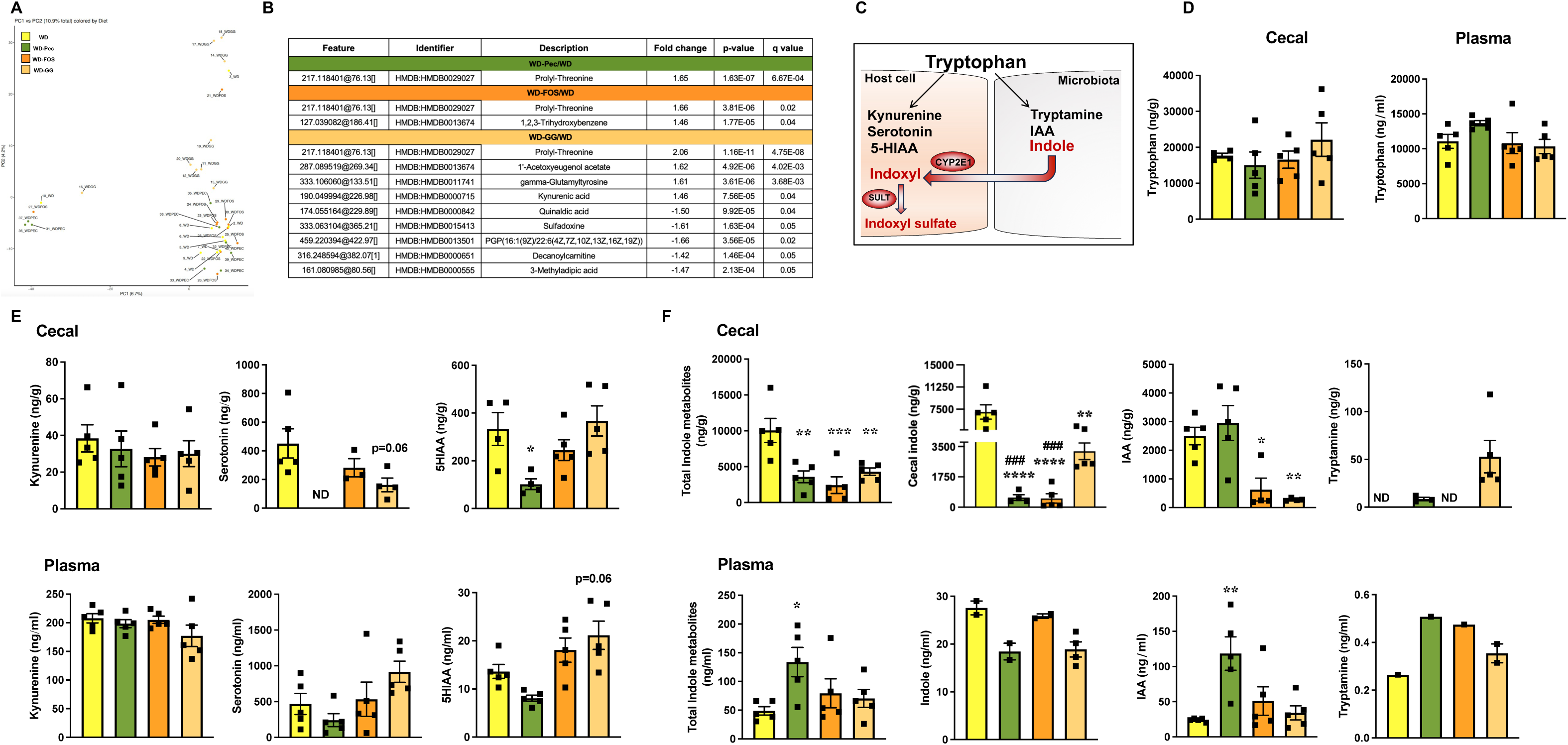
Dietary fiber supplementation contributes to subtle changes in the plasma metabolome induced by the Western diet. A) PCA plot illustrating the distribution of samples based on their metabolomic profile. B) Significantly altered metabolites for each pair-wise comparison. Positive fold change indicates a higher value in WD-FF, and negative fold change indicates a higher value in the WD group. C) Schematic of tryptophan catabolism pathways. D) Cecal and plasma tryptophan levels. Cecal and plasma levels of tryptophan metabolites E) host cell-generated -kynurenine, serotonin, 5-HIAA, and F) gut microbiome-generated-total metabolites (indole, IAA, tryptamine), indole, IAA, and tryptamine. n=9-10 per group. Differential expression analysis was performed using limma package. Adjusted P values (Q values) were calculated using the Benjamini-Hochberg false discovery rate correction and considered significant with a Q value <0.05 for A. Data analyzed by 1-way ANOVA with Sidak post-hoc tests for D-F. PCA, principal coordinate analysis; 5-HIAA, 5-hydroxy indoleacetic acid; IAA-indoleacetic acid. *P < 0.05; **P < 0.005; ***P < 0.001; ****P < 0.0001; ^###^P < 0.001. * WD vs WD-FF; ^#^ WD-GG vs WD-Pec or WD-FOS.

A small portion of dietary tryptophan is metabolized by host cells (via the kynurenine and serotonin pathways), and the remainder is metabolized by gut bacteria into indole and its derivatives, which then reach the liver via portal circulation (Figure 3C) ^25, 62^. Therefore, utilizing targeted LC-MS/MS, we next determined the levels of host and gut microbial tryptophan metabolites in cecal contents and plasma of the experimental mice. All diet groups had similar levels of plasma and cecal tryptophan (Figure 3D and Suppl. Figure 4C). Obesogenic diets shift tryptophan metabolism toward kynurenine production in host cells ^63, 64^. However, cecal and plasma kynurenine levels were not different between WD and CD (Suppl. Figure 4D) or WD and WD-FF groups (Figure 3E), likely due to differences in the composition of the experimental diets used (kcal: 40% fat, 28% sucrose vs 45% fat, 17% sucrose ^64^ and 60% fat ^63^). Cecal and plasma serotonin levels, another host-derived tryptophan metabolite, were significantly higher in the WD-fed compared to CD-fed mice (Suppl. Figure 4D). While in WD-FF groups, cecal serotonin was either undetected (WD-Pec) or trended lower (WD-GG) as compared to WD-fed mice (Figure 3E). A similar profile was observed for 5-hydroxyindoleacetic acid (5-HIAA) levels, a metabolite derived from serotonin. While WD has been previously shown to increase serotonin synthesis in mice, gut microbiome-dependent changes and inflammation can also contribute to these effects ^65^. Analysis of the concentration of gut bacteria-derived indole metabolites revealed that WD-fed mice had significantly higher levels of total cecal indole metabolites as compared to mice fed WD-FF (Figure 3F) or CD (Suppl. Figure 4E). When analyzed individually, markedly higher cecal indole levels in WD-fed mice were a major contributor to total indole metabolites. Cecal levels of other gut microbiota-derived tryptophan metabolites like indole acetic acid (IAA) and tryptamine varied differently among diet groups. Compared to WD, cecal IAA levels were similar in CD and WD-Pec groups but significantly lower in WD-FOS and WD-GG groups (Figure 3F and Suppl. Figure 4E). Tryptamine was not detected in WD and WD-FOS mice. In the plasma, total indole metabolite levels were significantly higher in WD-Pec mice and showed a trend toward higher levels in WD-GG, WD-FOS, and CD mice, with IAA being the primary contributor to this increase (Figure 3F and Suppl. Figure 4E).

### Enrichment of Western Diet with Fermentable Fiber Alleviates Gut Inflammation

Obesity and metabolic syndrome in human cohorts and murine models are associated with reduced cecal levels of indole and its derivatives ^63, 64^. In the gut, these metabolites suppress proinflammatory mucosal changes and leaky gut by activating the arylhydrocarbon receptor (AHR) and/or the pregnane X receptor (PXR), thereby restraining the development of systemic insulin resistance ^66, 67^. Thus, we first analyzed ileal mRNA expression of the indole metabolite receptors *Ahr* and *Pxr,* as well as their downstream targets, *Cyp1a1* and *Cyp3a11*, respectively, and found no significant differences among the diet groups (Suppl. Figure 5A). Our WD-fed mice had higher cecal indole levels compared to WD-FF mice (Figure 3F). It is notable that the major indole oxidizing isoform Cytochrome P 450 (CYP) 2E1 is also expressed in the intestine ^68, 69^, where, along with sulfotransferase (SULT)1A1, it participates in the conversion of indole to indoxyl sulfate ^70^. Thus, we next examined mRNA expression of *Cyp2e1* and *Sult1a1* in the ileum and distal colon. In the ileum, *Cyp2e1* expression was not statistically different between the WD and WD-FF groups; however, *Sult1a1* expression was significantly reduced in the Pec and FOS groups and showed a trend toward lower expression in the GG group (Figure 5A). Contrarily, in the distal colon, *Cyp2e1* expression was lower in all FF groups, reaching significance in the FOS and GG groups, while *Sult1a1* remained unchanged (Figure 5B). In CD-fed mice, *Cyp2e1* and *Sult1a1* expression was lower as compared to WD-fed mice, but not statistically different (Suppl. Figure 5B-C). Gut *Cyp2e1* induction in response to stresses like alcohol and fructose promotes oxidative stress and barrier damage ^69, 71^. Accordingly, the pore-forming tight junction protein claudin-2 (*Cldn 2*), which shows strong expression in the inflamed gut ^72^, was significantly higher in the ileum of WD-fed mice compared to WD-FF (Figure 4C) and CD-fed mice (Suppl. Figure 5D). Strikingly, expression of barrier-forming claudin, claudin-4 (*Cldn 4*), was also significantly higher in WD distal colon compared to CD (Suppl. Figure 5E) and WD-GG and WD-FOS fed mice (Figure 4D). This could be a compensatory response, as claudin-4 has been recently shown to selectively disrupt the membrane network of pore-forming claudins, such as claudin-2 ^73^. Expression of NADPH oxidase, *Nox4*, which generates reactive oxygen species and is upregulated in the inflamed gut^74^, trended low in the distal colon of WD-FF mice and reached significance in the WD-Pec group (Figure 4E). However, no difference was noted between CD and WD-fed mice (data not shown). WD induces a pro-inflammatory state in the gut ^54, 72, 75^ , as suggested by trending higher expression of cyclooxygenase-2 (*Cox2*) and *Ccl2* (Suppl. Figure 5D) in the ileum and antimicrobial peptide lipocalin-2 (*Lcn2*) (Suppl. Figure 5E) in the distal colon of WD-fed mice compared to CD-fed mice. Fiber supplementation of WD partially reversed these effects and significantly reduced the levels of *Il6* and *Ccl2* in the ileum, with *Lcn2* levels declining only in the WD-GG group (Figure 4F). *Il6* expression was also significantly reduced in the distal colon of all WD-FF groups (Figure 4G).

**Figure 4.**
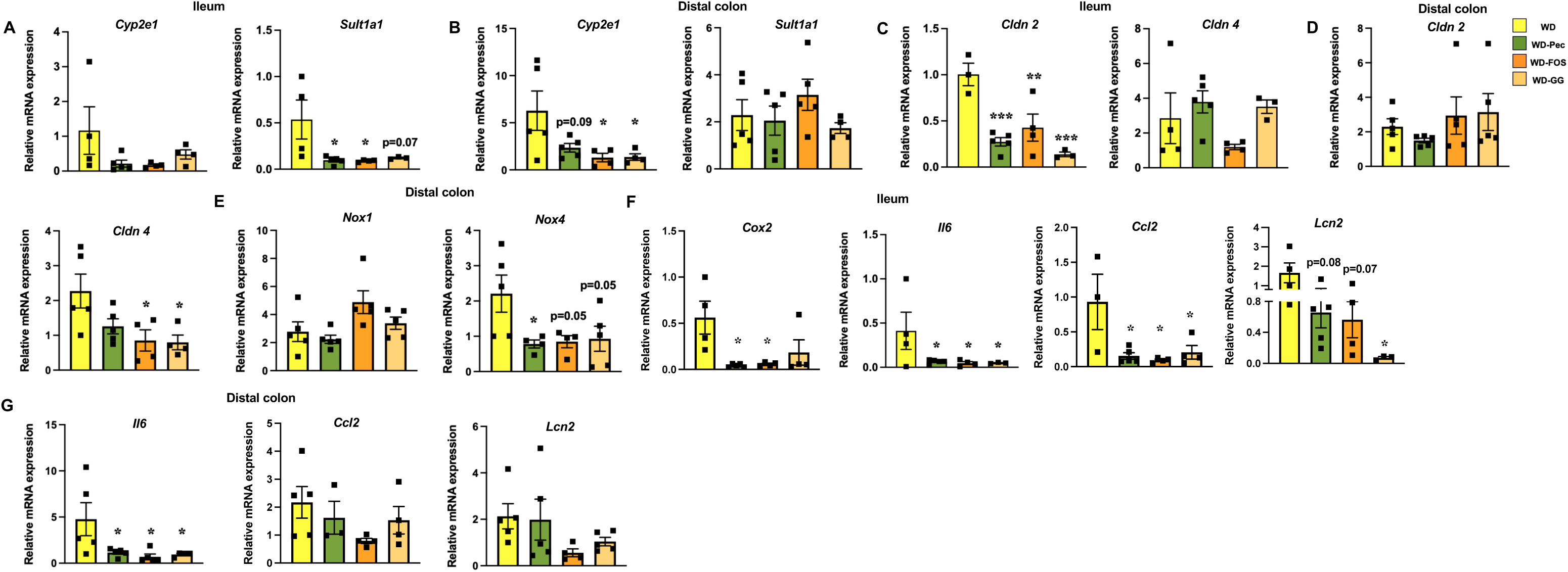
Dietary fiber supplementation mitigates gut inflammation induced by a Western diet. Levels of *Cyp2e1* and *Sult1a1* mRNA in A) the ileum and B) the distal colon. Levels of barrier genes (*Cldn 2* and *Cldn 4*) mRNA in C) the ileum and D) the distal colon. Levels of oxidative stress genes (*Nox1* and *Nox4*) mRNA in E) the distal colon and *Cox2* in F) the ileum. Levels of inflammatory genes (*Il6*, *Ccl2*, and *Lcn2*) mRNA in F) the ileum and G) the distal colon. n=3-5, Data are mean ± SEM analyzed by 1-way analysis of variance with Sidak post hoc analysis. *Cyp2e1*, cytochrome P 450 (CYP) family 2 subfamily E member 1; *Cox2*, cyclooxygenase-2; *Sult1a1*, sulfotransferase family 1A member 1; *Cldn 2*, claudin 2; *Cldn 4*, claudin 4; *Nox1*, NADPH oxidase 1; *Nox4*, NADPH oxidase 4; *Il6*, interleukin-6; *Ccl2*, C-C motif chemokine ligand 2; *Lcn2*, lipocalin 2. *P < 0.05; **P < 0.005; ***P < 0.001.

**Figure 5.**
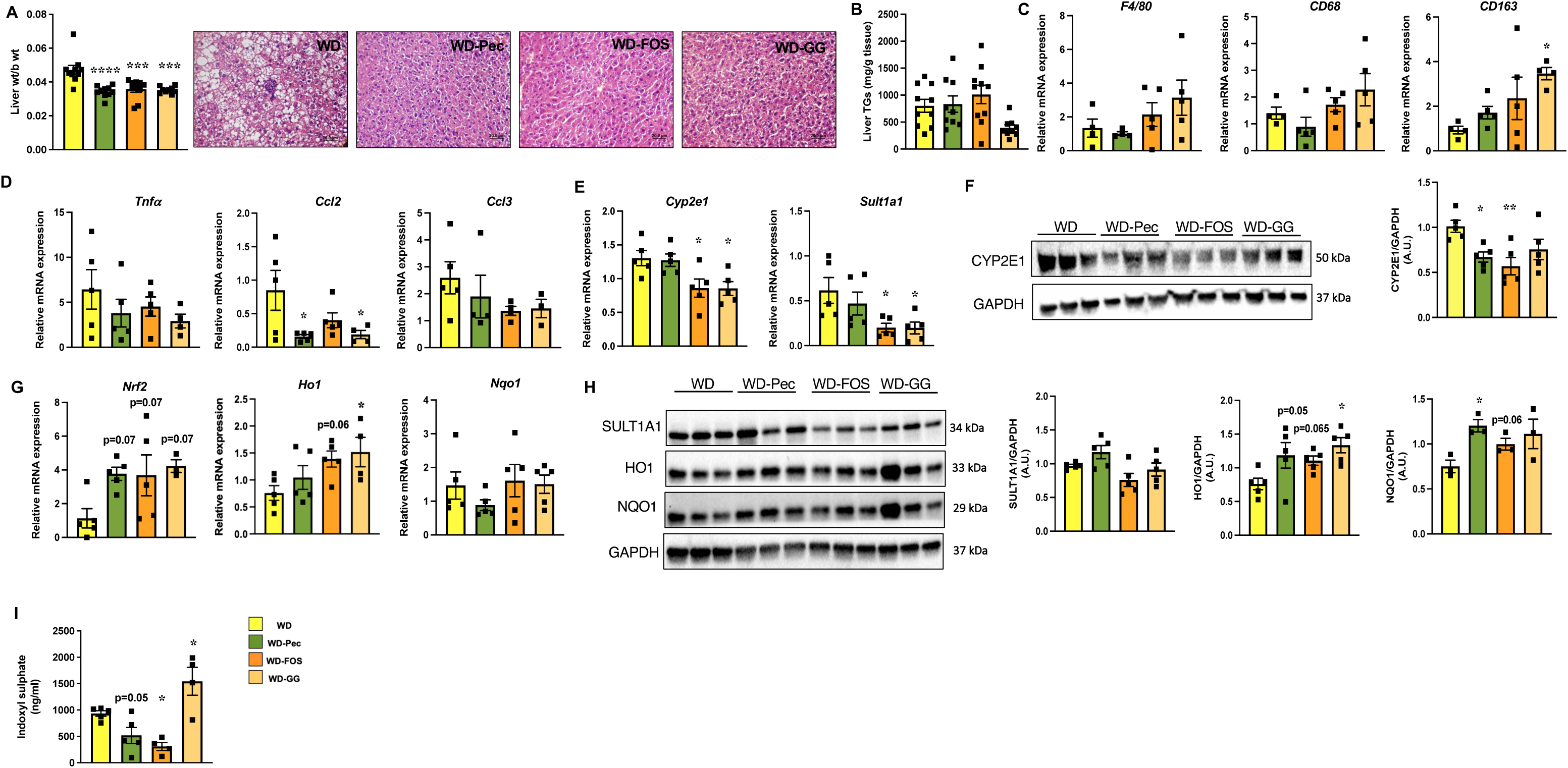
Dietary fiber supplementation alleviates Western diet-induced hepatic *Cyp2e1*, inflammation, and oxidative stress. A) Liver weights, histological analyses of liver tissue by H&E staining, and NAFLD activity score (NAS) score consisting of steatosis, ballooning, and inflammation. B) Liver triglycerides. C) and D) mRNA expression of immune cell markers, *F4/80*, *CD68*, *CD163*, and cytokine genes, *TNF*𝘢, *Ccl2*, and *Ccl3*, respectively. Expression of E) *Cyp2e1* and *Sult1a1* mRNA and F-H) CYP2E1, SULT1A1, HO-1, NQO1 proteins with corresponding densitometric analysis. G) mRNA expression of *Nrf2*, *Ho1*, and *Nqo1*. I) Plasma levels of indoxyl sulfate. Data are mean ± SEM; n=9-10 per group for A sub-panel on liver weights and B; n=3-5 per group for remaining sub-panels. Data were analyzed by 1-way analysis of variance with Sidak post hoc analysis. NAFLD, nonalcoholic fatty liver disease; *F4/80*, EGF-like module-containing mucin-like hormone receptor-like 1; *CD68*, cluster of differentiation 68; *CD163*, cluster of differentiation 163; *TNF*𝘢, tumor necrosis factor 𝘢; *Ccl2*, C-C motif chemokine ligand 2; *Ccl3*, C-C motif chemokine ligand 3; *Cyp2e1*, cytochrome P 450 (CYP) family 2 subfamily E member 1; *Sult1a1*, sulfotransferase family 1A member 1; *Nrf2*, nuclear factor erythroid 2-related factor 2; *Ho-1*, heme oxygenase-1; *Nqo1*, NAD(P)H quinone oxidoreductase 1. *P < 0.05; **P < 0.005; ***P < 0.001; ****P < 0.0001.

### Fermentable Fiber Alleviates Western Diet-Induced Hepatic Cyp2e1, Inflammation, and Oxidative Stress

Low-fiber diets are associated with an increased risk of liver inflammation and steatosis ^76^. While our CD-fed mice had similar liver weights and hepatic TG content as WD-fed mice (Suppl. Figure 6A and C), the histological analysis via H&E staining revealed higher fat accumulation in WD livers (Suppl. Figure 6B). Upon analysis of de novo lipogenesis (DNL) pathway genes, only carbohydrate-responsive element–binding protein, *Chrebp*α, and *Chrepb*β expression was higher in CD-fed compared to WD-fed livers (Suppl. Figure 6D). In view of CD-fed mice exhibiting higher insulin sensitivity and glucose tolerance compared to WD-fed mice (Suppl. Figure 2J and K), there appears to be a dissociation between hepatic TGs and insulin resistance noted in mice with overexpression of CHREBP ^77^. On the other hand, the livers from WD-FF-fed mice weighed significantly less, and their histological analyses revealed reduced fat accumulation compared to the WD group (Figure 5A). Interestingly, as observed in CD-fed mice, liver TGs were not significantly different among the diet groups (Figure 5B). FFs impact hepatic lipid metabolism by reducing DNL and concomitantly increasing fatty acid oxidation ^78^. Indeed, several lipid metabolism genes showed similar differential regulation in WD-FF groups, with the most significant changes noted with increased *Chrebp*α, *Chrepb*β, and *Ldlr* (low density lipoprotein receptor) in WD-Pec, reduced *Fasn* (fatty acid synthase) in WD-GG, reduced *Gpat* (glycerol-3-phosphate acyl transferase) in WD-FOS and WD-GG, and increased *Cpt2* (mitochondrial carnitine palmitoyltransferase II) in WD-Pec and WD-FOS groups (Suppl. Figure 7). Thus, in WD-FF mice, FFs appear to improve hepatic lipid handling and turnover. Furthermore, WD-fed mice exhibited higher inflammation marked by increased expression of *Cd68* and *Vsig4* (V-set and immunoglobulin domain containing 4) markers for Kupffer cell activation compared to CD-fed mice (Suppl. Figure 6E). Overall, immune cell activation markers were not different between the WD and WD-FF groups, except *Cd163*, a marker expressed by anti-inflammatory macrophages which is also inversely associated with hepatic steatosis ^79^, where it was significantly higher in the WD-GG group (Figure 5C). Expression of *Ccl2*, a chemokine associated with hepatic damage in obesity related diseases ^80^, was significantly reduced in WD-Pec and WD-GG groups compared to WD-fed mice (Figure 5D).

As noted earlier, WD-FF groups exhibited significantly reduced fecal levels of indole (Figure 3F). In the liver, indole is 3-hydroxylated and converted to indoxyl by the activity of hepatic CYP2E1, and then O-sulfated by SULT1A1, yielding indoxyl sulfate (Figure 3C)^81^. Indole and its derivatives, in general, are beneficial, except for indoxyl sulfate, which becomes a uremic toxin at high concentrations ^67^. To explore this indole-specific pathway in our experimental mice, we analyzed the expression of CYP2E1 and SULT1A1 in the liver. No differences were noted in the mRNA or protein expression between CD and WD-fed mice (Suppl. Figure 6F and G). mRNA levels of *Cyp2e1* and *Sult1a1* were significantly downregulated in WD-FOS and WD-GG groups (Figure 5E), while reduced protein levels of CYP2E1 were noted in WD-PEC and WD-FOS groups (Figure 5F). Hepatic CYP2E1 has been shown to directly cause fatty liver disease by increasing oxidative stress ^71^. Conversely, the induction of *Nrf2* (nuclear factor erythroid 2-related factor 2) and its downstream target antioxidant enzymes, *Ho-1* (heme oxygenase-1) and *Nqo1* (NAD(P)H: quinone oxidoreductase 1), is a key adaptive response against CYP2E1 ^82, 83^. Accordingly, as compared to the WD group, mRNA expression of *Nrf2* trended high in the livers of WD-FF mice, and that of *Ho-1* was significantly high in the WD-GG group (Figure 5G). A corresponding significant increase was also noted in the protein levels of HO-1 in the WD-GG and NQO-1 in the WD-Pec group (Figure 5H). Finally, the plasma concentration of the product of CYP2E1 and SULT1A1 activity on indole, indoxyl sulfate, was observed to be lower, modestly in WD-PEC and significantly in WD-FOS groups, compared to the WD group (Figure 5I). In contrast, WD-GG showed a significantly high plasma concentration of indoxyl sulfate (Figure 5I). Consequently, expression of the xenobiotic receptor, *Pxr*, and of its downstream target detoxifying enzyme *Cyp3a11* ^84, 85^, was significantly high in the liver of the WD-GG group (Suppl. Figure 8A). Lastly, plasma levels of total and individual SCFAs, which more directly relate to metabolic health ^86^, were higher in WD-FF groups than in WD-fed mice, reaching significance in the WD-FOS group (Figure 6A). CD and WD-fed mice had similar plasma indoxyl sulfate (Suppl. Figure 6H). Although total SCFA and acetate levels were similar between the CD- and WD-fed groups, propionate levels were significantly higher in CD-fed mice (Suppl. Figure 6I).

**Figure 6.**
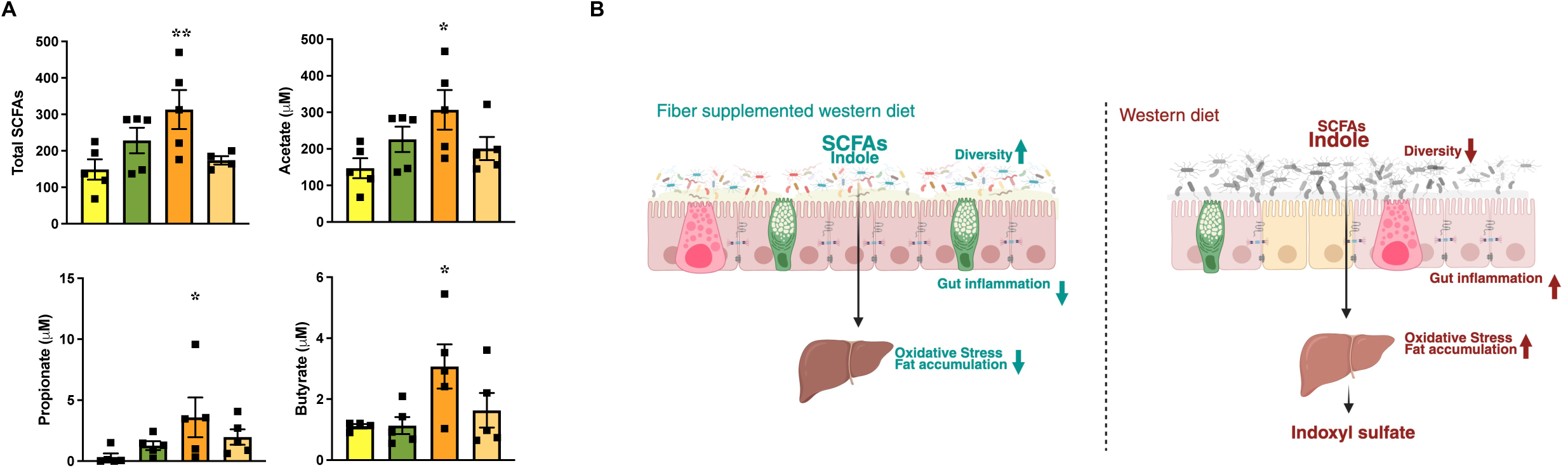
Plasma short-chain fatty acids following dietary fiber supplementation. A) Plasma total SCFAs and individual SCFAs (acetate, butyrate, and propionate) levels. B) Fiber supplementation of the Western diet restores gut microbial diversity and levels of its metabolites, SCFAs, and indole, leading to reduced gut inflammation, hepatic oxidative stress, and fat accumulation. Conversely, the typical Western diet disrupts gut microbial diversity, leading to a reduction in SCFA production and an increase in indole levels, which contribute to the formation of indoxyl sulfate in the liver. Data are mean ± SEM; n=4-5, 1-way analysis of variance with Sidak post hoc analysis. SCFAs, short-chain fatty acids. *P < 0.05; **P < 0.005.

## Discussion

Our study highlights two key findings: a recognized finding, that FFs are anti-obesogenic ^17^, and a second, unique finding that FFs can reduce indoxyl sulfate levels through modulation of hepatic CYP2E1 and SULT1A1. Both point towards the emerging potential of dietary fiber as a therapeutic modality against NCDs.

A large body of evidence supports the role of WD consumption in increasing visceral fat, intestinal barrier permeability, and hepatic fat accumulation in the context of systemic inflammation and the propagation of obesity-associated metabolic disorders. Currently, therapies targeting the enteroendocrine pathways are most effective for weight loss and reducing these adverse effects ^17^. However, their use is limited due to affordability, gastrointestinal tolerance issues, and insufficient knowledge about their long-term health effects ^17^. Thus, diet modulation appears to be a cost-effective and generally well-tolerated option for weight loss. In this regard, various dietary fibers have been investigated for their metabolic and weight loss benefits in preclinical models and clinical cohorts^17, 18, 78^. FFs affect metabolic health through the combined effects of their physicochemical properties-solubility, viscosity, and fermentability-which collectively modulate digestion, lipid metabolism, glycemic control, energy expenditure, and gastrointestinal motility. We recapitulated several documented beneficial effects of dietary FF on dysregulated metabolic profiles induced by WD (Figure 7B). However, we did not observe marked differences in energy expenditure and food intake, suggesting reduced lipid absorption and, consequently, storage, along with the metabolic effects of SCFAs as the primary contributors to the lean phenotype of our WD-FF mice. Notably, FOS, a soluble but non-viscous FF, exerted the least weight loss benefit compared to gel-forming GG and Pec ^87^. While our preclinical WD-induced obesity model demonstrates weight loss and metabolic benefits from the consumption of isolated, purified fiber, translating these effects to the clinic is not straightforward. Variable dietary choices, diet adherence, fiber tolerance, and baseline gut microbial differences can all impact the effects of fiber. For example, inflammatory responses in IBD patients induced by FF are suggested to be due to the absence of key fermenter strains ^88^, and the selection of specific FF responders in type 2 diabetic subjects can promote glycemic control ^9^. A key approach will be to use retrograde translational or humanized mouse models ^89, 90^ with the gut microbiome of obese individuals to identify the similar weight loss potential of FFs in the presence of persistent obesogenic stress.

Fermentation of the fibers by the gut microbiome yields a multitude of metabolites and selects for specific gut bacterial strains through cross-feeding. In our cohorts, we did not observe specific spikes in gut bacterial families earlier shown for WD versus FF, such as *Enterobacteriaceae* versus *Prevotellaceae* ^18^. In fact, FF groups exhibited a greater proportion of *Enterobacteriaceae*. *Proteobacteria* have been shown to correlate positively with obesity ^7, 8^, and FF consumption has been found to reduce their levels ^91^. Several reasons can account for the increase in these facultative anaerobes in WD-FF mice. This may be an outcome of complex community interactions in the presence of persistent metabolic stress, as the development of a community is influenced by the interactions between the organisms present and their resource environment ^92^. WD induces an inflamed gut characterized by increased reactive oxygen and nitrogen species, a prominent factor promoting the growth of *Enterobacteriaceae* ^8^. FF consumption would further trigger complex metabolic interactions among the gut microbial community, whereby by-products of community members can indirectly stimulate the growth of other members ^93^. For example, the fermentation of complex carbohydrates by members of *Bacteroidetes* can produce simpler monosaccharides and dicarboxylic acids, which can promote the growth of *Enterobacteriaceae* ^8, 49, 52^. Notably, our WD-FF-fed mice exhibited a significant expansion of *Bacteroidetes* members (Figure 2C). Importantly, the expansion of *Enterobacteriaceae* members did not overwhelm the system negatively. Interestingly, a similar increase in *Proteobacteria* and members of *Enterobacteriaceae* was observed in CD-fed mice (Suppl. Figure 3F), where a well-studied shift in the pH and oxygen availability could likely favor the facultative anaerobes ^8^. Thus, our data indicate that the dominance of a single purified fermentable fiber in a western-style diet or an absolute lack thereof can select for specific microbial communities over time. Understanding if this potentially impacts the course of secondary complications of NCDs or the occurrence of new insults, such as colitis ^49, 94^, requires further research.

These gut microbial profiles may also partially support the limited alterations we observed in the plasma metabolome of our mouse cohorts. However, FFs were able to select SCFA producers. In addition to increased plasma levels of SCFAs, our mouse cohorts revealed altered tryptophan metabolites. Higher cecal indole and reduced plasma indole derivatives in WD-fed mice, and the opposite trend of reduced cecal indoles and higher plasma indole derivatives in WD-FF mice were observed. FFs have recently been demonstrated to correlate with reduced indole production but increased indole derivatives, such as indole propionic acid (IPA), in both rodent ^61^ and human cohorts ^9, 95–97^. Sinha et al. ^61^ showed that cross-feeding pathways initiated by fiber degraders like *B. thetaiotaomicron* shift available tryptophan from indole producers to Stickland fermenters (that are capable of coupling oxidation and reduction of amino acids) to produce alternate tryptophan derivatives like IAA and IPA. Additionally, several *Bifidobacterium* and *Lactobacillaceae* species can directly convert tryptophan to metabolites other than indole ^59, 98–100^. While the possibility of these diverse pathways occurring in our cohort exists and needs further exploration, increased proteolytic fermentation in WD-fed mice could also contribute to higher cecal indole. In fact, we observed significantly increased cecal levels of isobutyric acid and valeric acid, products of proteolytic fermentation ^101^ in WD-fed mice (Suppl. Figure 8B). Importantly, our panel for targeted metabolomics of tryptophan metabolites did not include IPA and indole lactic acid. Still, we observed higher plasma IAA in WD-FF mice. IAA demonstrates protective effects against metabolic disorders associated with obesity and diabetes ^102^. In rodent models of non-alcoholic liver disease and diet-induced obesity, IAA supplementation has been shown to reduce oxidative stress, inflammation, and insulin resistance, as well as improve lipid metabolism^63, 103^. Thus, in addition to the direct metabolic effects of dietary FF, altered tryptophan metabolites, such as IAA, could contribute to the protected phenotype observed in WD-FF mice.

Higher plasma indoxyl sulfate levels were observed in WD-fed mice. Complementing this, higher CYP2E1, oxidative stress, and inflammation were observed in the livers of WD-fed mice. Compared to other groups, WD-GG mice exhibited higher plasma indoxyl sulfate levels and a compensatory increase in hepatic expression of *Pxr* and *Cyp3a11*. The production of indoxyl sulfate is reportedly independent of genetic variations in CYP2E1 and SULT1A1, but rather depends on gut microbiome variability ^104^. IAA and indoxyl sulfate production can also be affected by gut transit time ^104^. It is possible that higher transit times, supported by a highly viscous guar gum, contribute to higher cecal indoles in WD-GG compared to WD-Pec and WD-FOS mice (Figure 3F), which could ultimately lead to higher plasma indoxyl sulfate levels in this group. Thus, a complex interplay between diet and gut microbiome likely determines indoxyl sulfate production. Although we did not examine kidney function in our mouse cohort, our results suggest that certain dietary FFs may be used to mitigate the risk of chronic kidney disease occurring as a comorbidity with metabolic disorders. However, first, it will be crucial to determine in preclinical models the minimum effective dose, type, and duration of fiber treatment, as well as the enrichment of specific gut bacterial pathways, and whether sexual dimorphism exists in fiber-induced suppression of indole production, given that our study used only males. Regardless, understanding these complex effects could lead to novel diet-based therapies that target NCDs.

## Acknowledgements

This publication includes data generated at the UC San Diego IGM Genomics Center using an Illumina NovaSeq 6000, which was purchased with funding from a National Institutes of Health SIG grant (#S10 OD026929). Bioinformatics analysis in the project described was performed by the UIC Research Informatics Core, supported in part by NCATS through Grant UM1TR005438.

## Abbreviations

Ahr: aryl hydrocarbon receptor
ANOVA: analysis of variance
Acc1: acetyl-CoA carboxylase
Acly: ATP citrate lyase
AUC: area under the curve
Ccl2: C-C motif chemokine ligand 2
Ccl3: C-C motif chemokine ligand 3
Cd36: cluster of differentiation 36
CD68: cluster of differentiation 68
CD163: cluster of differentiation 163
Chrebpα: carbohydrate-responsive element-binding protein isoform α
Chrebpβ: carbohydrate-responsive element-binding protein isoform β
Cldn4: claudin 4
Cldn 2: claudin 2
Cox2: cyclooxygenase-2
Cyp2e1: cytochrome P 450 (CYP) family 2 subfamily E member 1
CD: control diet
Cpt2: carnitine Palmitoyltransferase II
Cyp3a11: cytochrome P 450 (CYP) family 3 subfamily A member 1
Fasn: fatty acid synthase
FF: fermentable fiber
F4/80: EGF-like module-containing mucin-like hormone receptor-like 1
FDR: false discovery rate
FOS: fructooligosaccahride
GG: guar gum
GM: gut microbiome
Gpat: glycerol-3-phosphate O-acyltransferase
5-HIAA: 5-hydroxy indoleacetic acid
Ho-1: heme oxygenase-1
HOMA-IR: homeostasis model assessment of insulin resistance
Il6: interleukin-6
IP-GTT: intraperitoneal glucose tolerance test
IAA: indoleacetic acid
ITT: insulin tolerance test
LC-MS: Liquid chromatography–mass spectrometry
Lcn2: lipocalin 2
Ldlr: low density lipoprotein receptor
NEFA: non-esterified fatty acid
PCA: principal coordinate analysis
Nox1: NADPH oxidase 1
Nox4: NADPH oxidase 4
Nrf2: nuclear factor erythroid 2-related factor 2
Nqo1: NAD(P)H quinone oxidoreductase 1
Pec: pectin
PERMANOVA: permutational multivariate analysis of variance
Pxr: pregnane X receptor
SCFA: short chain fatty acid
Srebp1c: sterol regulatory element binding transcription factor 1
Sult1a1: sulfotransferase family 1A member 1
Ho-1: heme oxygenase-1
Nqo1: NAD(P)H quinone oxidoreductase 1
WD: western diet.

## Supplementary Figure Legends

**Suppl. Figure 1.**
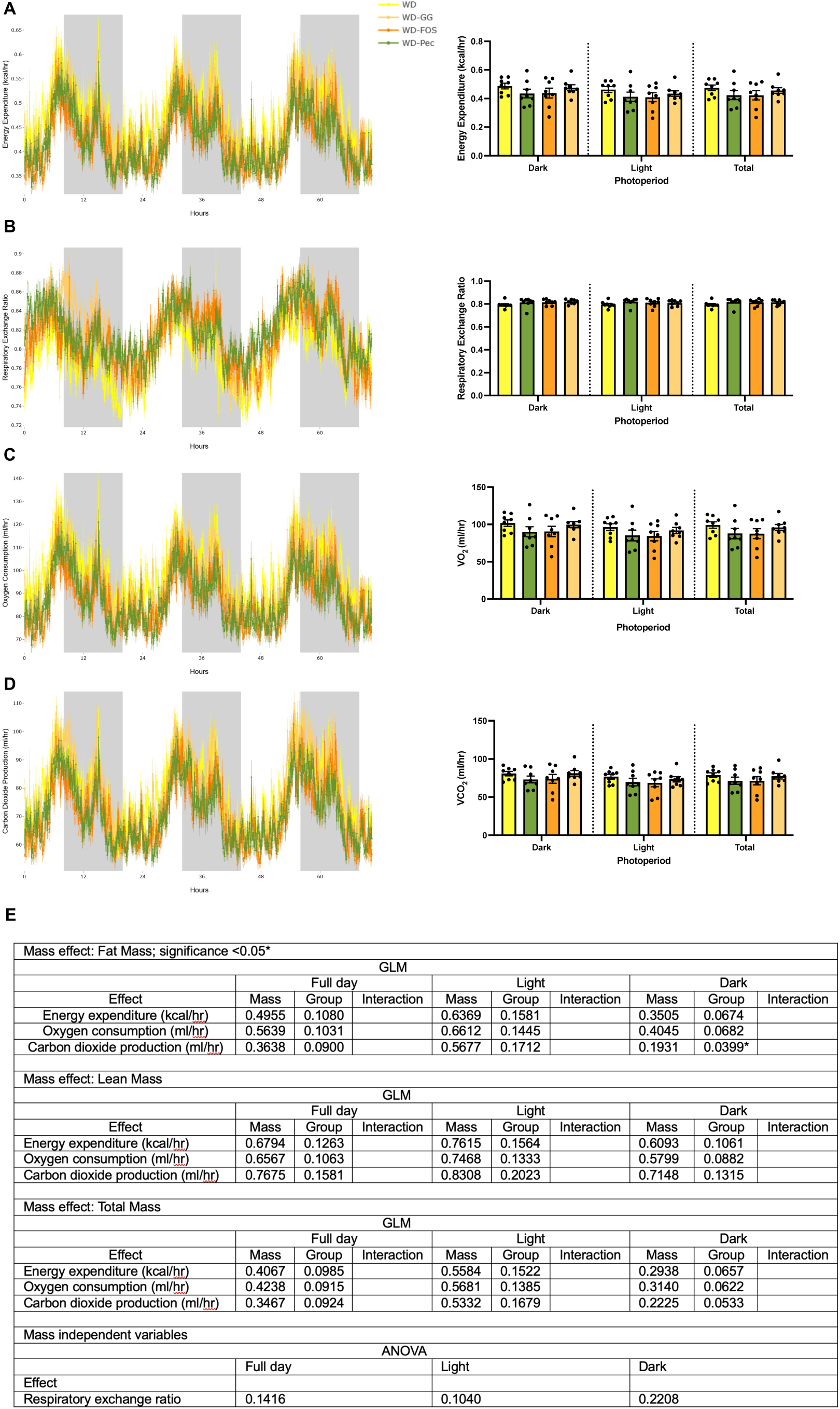
Dietary fiber supplementation drives subtle changes in energy expenditure or respiratory exchange ratio. A) Energy expenditure, B) Respiratory exchange ratio, C) Oxygen consumption (VO_2_), and D) Carbon dioxide production (VCO_2_) measured by indirect calorimetry, E) ANCOVA analyses run separately for dark, light, and whole day period. n=8 per group. GLM, generalized linear model.

**Suppl. Figure 2.**
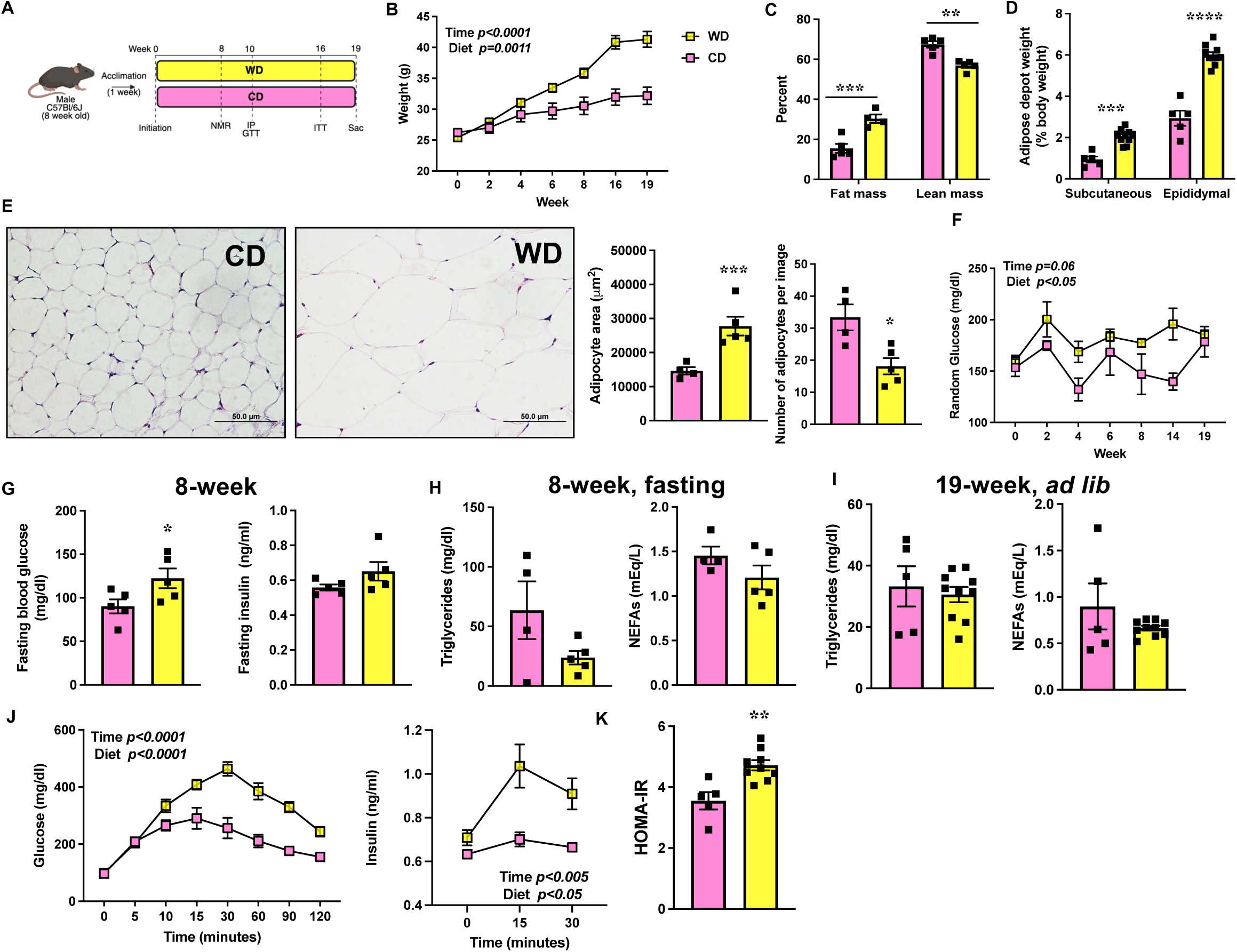
Phenotypic differences between mice on control and Western diets. A) Schematic of the experimental design. B) Body weight. C) Fat mass and lean mass at 12 weeks. D) Weights of subcutaneous and epididymal adipose tissues. E) Representative images of H & E-stained subcutaneous adipose tissue, average area of individual adipocytes, and number of adipocytes per image. F) Ad lib glucose levels measured throughout the study. G) 8-week fasting glucose and insulin levels. H) 8-week fasting and I) 19-week ad lib, triglycerides and NEFA levels. J) IP-GTT measured after 10 weeks on the diet and insulin levels during the first 30 minutes of the test. K) 19-week HOMA-IR. Data are mean ± SEM; n=5-10 per group for B, D, F, I, J, K; n=4-5 per group for C, E, G, H. Data were analyzed by 2-way ANOVA with Sidak post-hoc tests for B, F and J and without post hoc for C and D or Student’s two-tailed *t*-test for E, G, H, I and K. NEFA, non-esterified fatty acid; HOMA-IR, homeostasis model assessment of insulin resistance; IP-GTT, intraperitoneal glucose tolerance test; ANOVA, analysis of variance. *P < 0.05; **P < 0.005; ***P < 0.001; ****P < 0.0001.

**Suppl. Figure 3.**
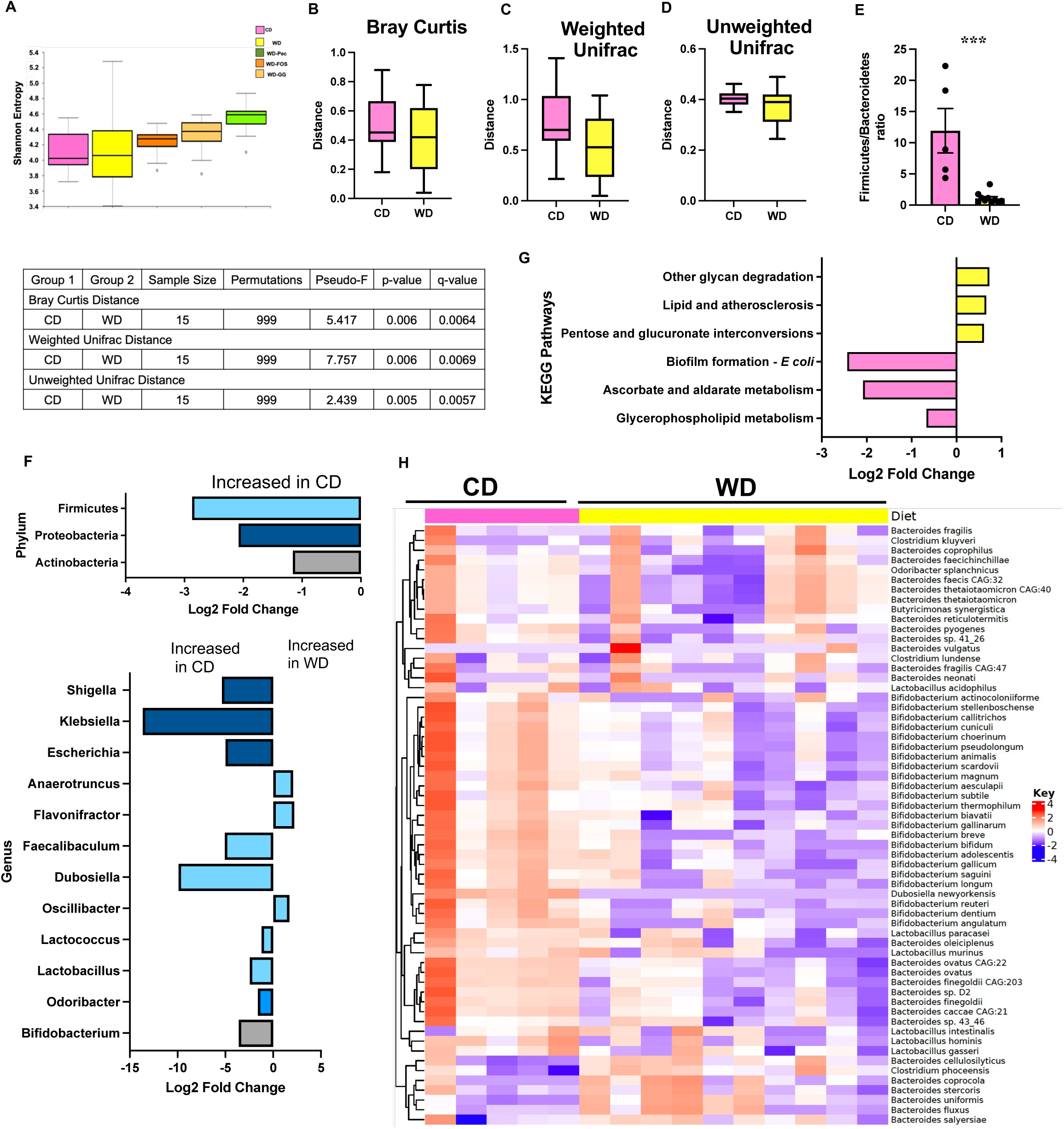
Alterations in the cecal gut microbiome of the different diet groups. A) Alpha diversity indicated by Shannon Entropy in different diet groups. Beta diversity: B) Bray-Curtis, C) Weighted, and D) Unweighted Unifrac distances. E) Firmicutes/Bacteroidetes percentage ratio presented as mean ± SEM. F) Bar plots displaying log2 fold changes at two taxonomic levels (phylum and genus) for pairwise comparisons between samples. G) Bar plots illustrating log2 fold changes for relevant KEGG pathways that were significantly enriched in the treatment groups and showed significance in the differential analyses. H) A heatmap showing species with significantly abundant changes from selected genera, constructed using log2-scaled and z-scored normalized abundances. Each row represents a species, while each column represents a sample.CD, n=5; WD, WD-FF, n=9-10 per group. Data analyzed by PERMANOVA for A and B, Student’s two-tailed *t*-test for E. For G and F, pairwise comparisons were performed using edgeR. Adjusted P values calculated using the Benjamini-Hochberg false discovery rate correction were considered significant with a Q value <0.05.

**Suppl. Figure 4.**
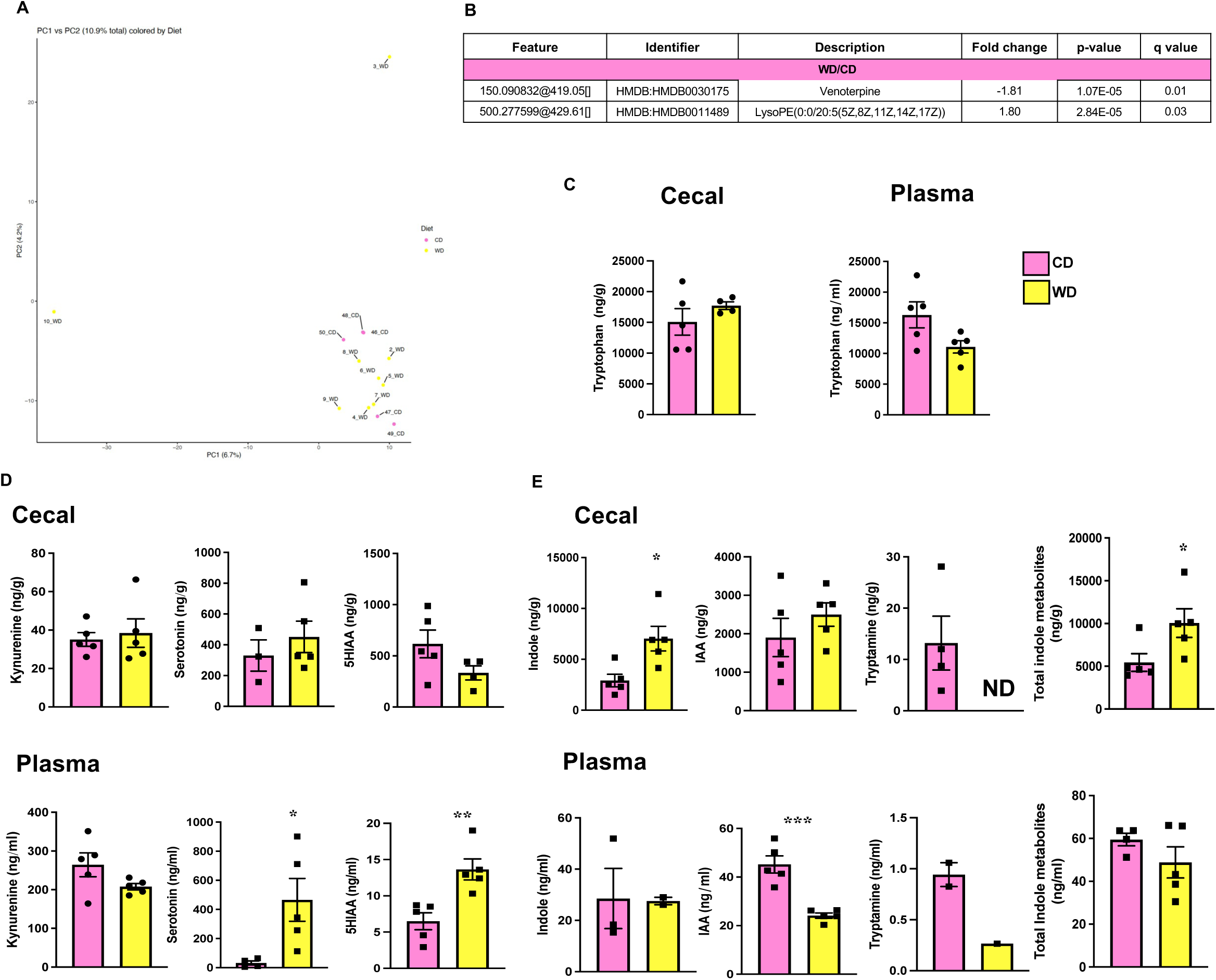
Untargeted and targeted metabolomic features in control and Western diet-fed mice. A) PCA plot. B) Significantly altered metabolites for WD-CD comparison with positive fold change signifying a higher value in WD, and negative fold change indicating a higher value in the CD group. C) Cecal and plasma tryptophan levels. D) and E) Cecal and plasma levels of tryptophan metabolites of host-derived (kynurenine, serotonin, 5-HIAA) and gut microbiome-generated (total or individual, indole, IAA, tryptamine). n=4-9 per group. Adjusted P-values (Q values) were calculated using the Benjamini-Hochberg false discovery rate correction, and were considered significant with a Q value <0.05 for A. For other panels, n=4-5 and data analyzed by Student’s two-tailed *t*-test. PCA, principal coordinate analysis; 5-HIAA, 5-hydroxy indoleacetic acid; IAA-indoleacetic acid. *P < 0.05; **P < 0.005; ***P < 0.001.

**Suppl. Figure 5.**
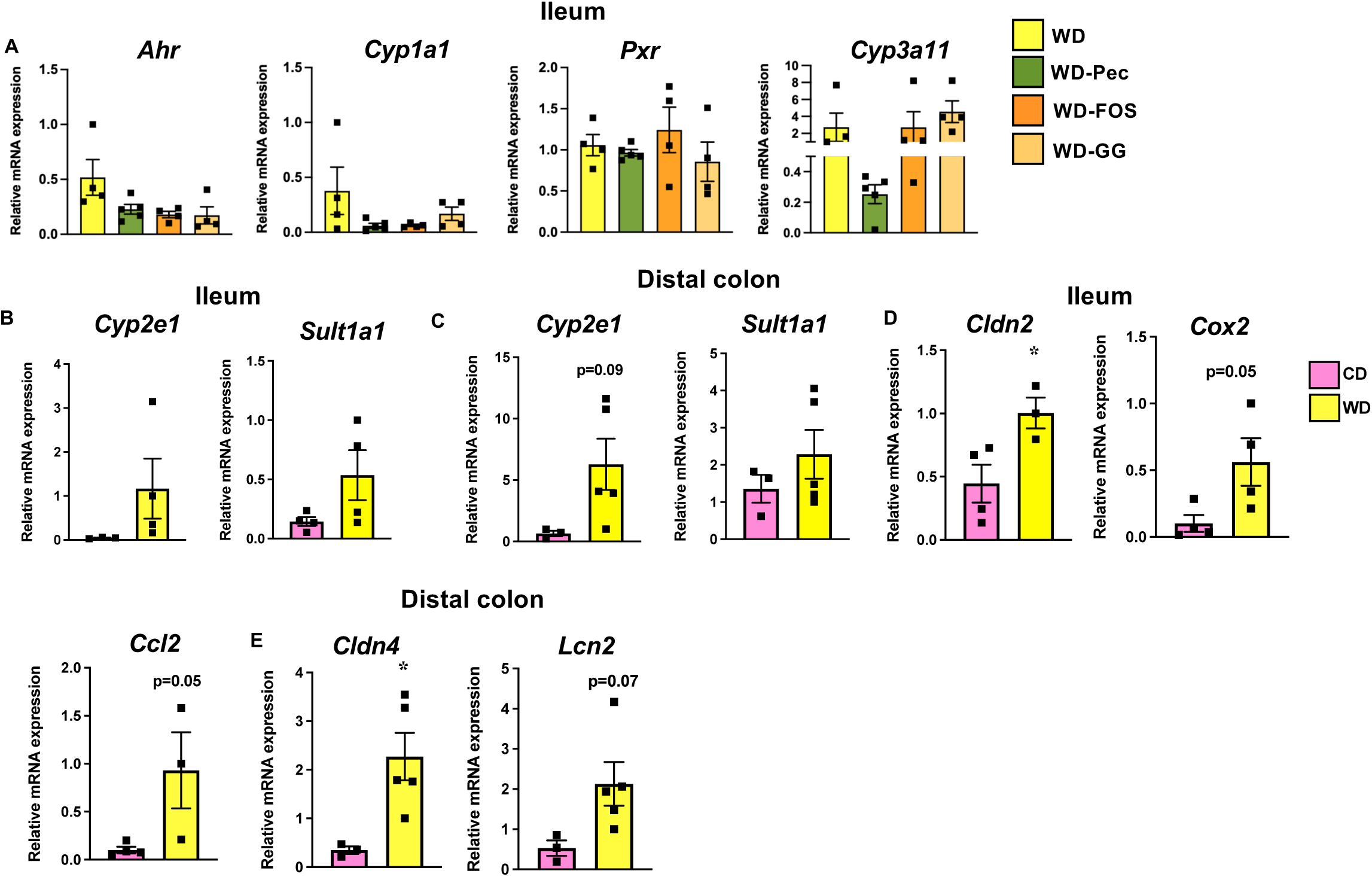
Expression of indole-specific, gut barrier, and inflammatory genes. A) Ileal mRNA expression of *Ahr*, *Cyp1a1*, *Pxr*, and *Cyp3a11* in WD and WD-FF mice. Levels of *Cyp2e1* and *Sult1a1* mRNA in B) the ileum and C) the distal colon of WD and CD fed mice. D) Levels of *Cldn 2* and *Cox2* mRNA in the ileum and of *Ccl2*, *Cldn 2* , and *Lcn2* in the distal colon. n=3-5, Data are mean ± SEM analyzed by 1-way analysis of variance with Sidak post hoc analysis for A and Student’s two-tailed *t*-test for other panels. *Ahr*, aryl hydrocarbon receptor; *Cyp1a1*, cytochrome P 450 (CYP) family 1 subfamily A member 1; *Cyp3a11*, cytochrome P 450 (CYP) family 3 subfamily A member 1; *Pxr*, pregnane X receptor; *Cyp2e1*, cytochrome P 450 (CYP) family 2 subfamily E member 1; *Sult1a1*, sulfotransferase family 1A member 1; *Cldn 2*, claudin 2; *Cldn 4*, claudin 4; *Ccl2*, C-C motif chemokine ligand 2; *Lcn2*, lipocalin 2. *P < 0.05.

**Suppl. Figure 6.**
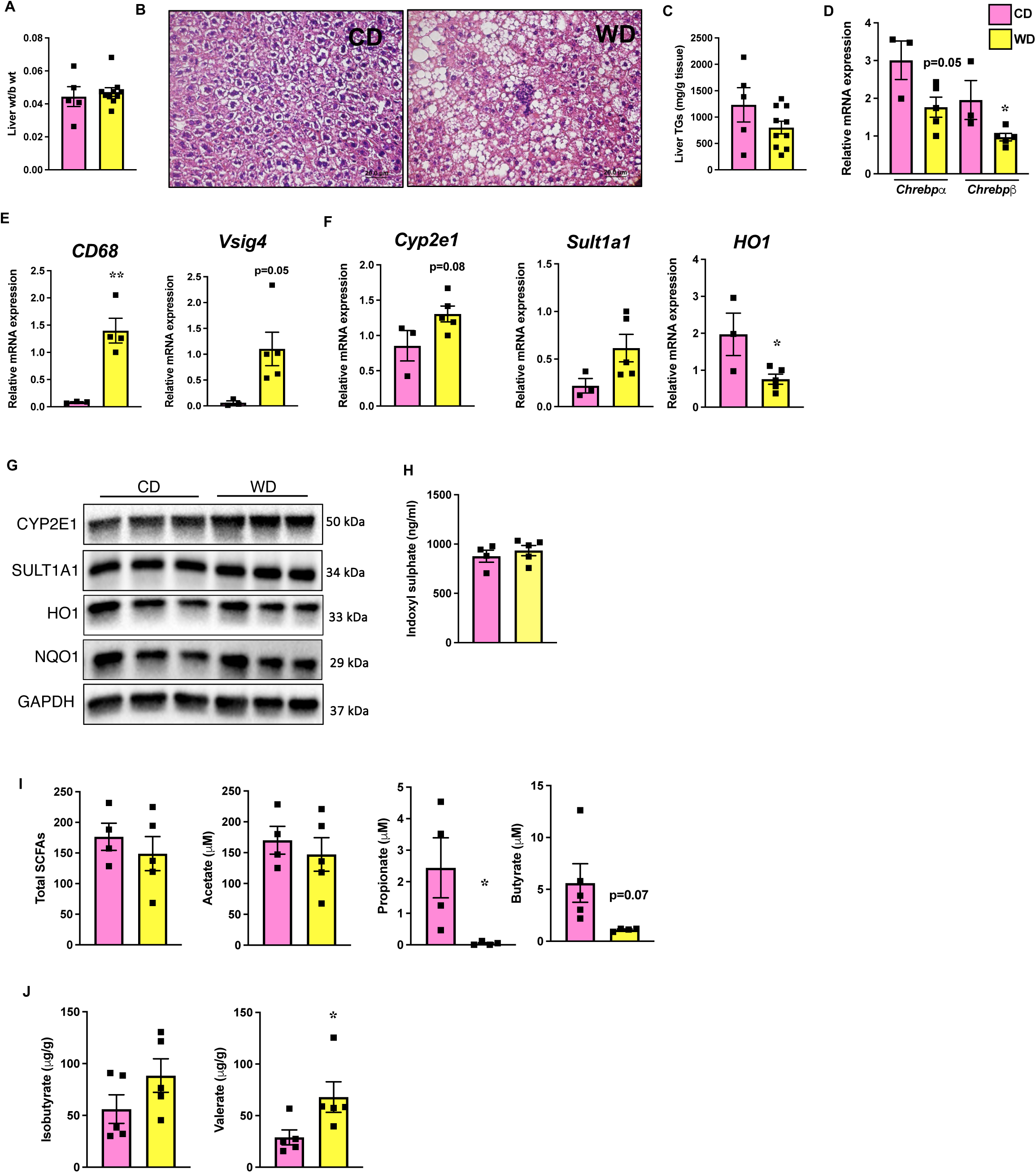
Liver inflammation and oxidative stress in WD and CD mice. A) Liver weights. B) histological analyses of liver tissue by H&E staining. C) Liver triglycerides. mRNA expression of D) lipid metabolic genes, E) immune cell markers, and F) *Cyp2e1*, *Sult1a1*, and *Ho-1*. G) Western blot of CYP2E1, SULT1A1, HO-1, and NQO1. H) Plasma levels of indoxyl sulfate. I) Plasma total and individual SCFAs. J) Cecal levels of isobutyrate and valerate. Data are mean ± SEM; n=3-10 per group for A and C, and n=3-5 for the remaining sub-panels. Data were analyzed by 1-way analysis of variance for D and Student’s two-tailed *t*-test for other panels. *CD68*, cluster of differentiation 68; *Vsig4*, v-set and immunoglobulin domain containing protein 4; TGs, triglycerides; *Cyp2e1*, cytochrome P 450 (CYP) family 2 subfamily E member 1; *Sult1a1*, sulfotransferase family 1A member 1; *Ho-1*, heme oxygenase-1; *Nqo1*, NAD(P)H quinone oxidoreductase 1. *P < 0.05.

**Suppl. Figure 7.**
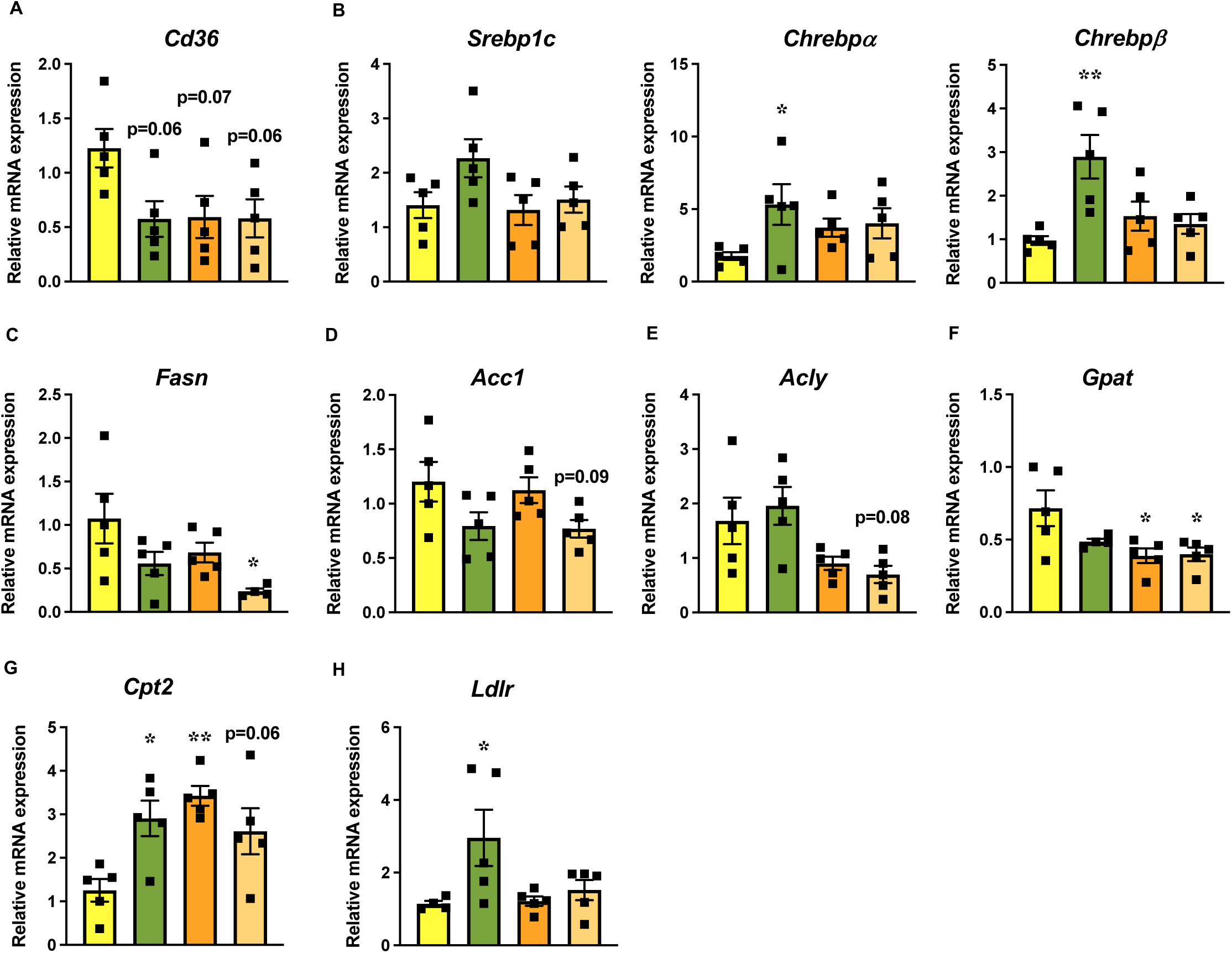
Expression of *de novo* lipogenesis and other lipid metabolism genes in the livers of WD and WD-FF fed mice. n=3-5 per group. Data are mean ± SEM; analyzed by 1-way analysis of variance with Sidak post hoc tests. *Cd36*, cluster of differentiation 36; *Srebp1c*, sterol regulatory element binding transcription factor 1; *Chrebp*α, carbohydrate-responsive element-binding protein isoform α; *Chrebp*β, carbohydrate-responsive element-binding protein isoform β; *Fasn*, fatty acid synthase; *Acc1*, acetyl-CoA carboxylase; *Acly*, ATP citrate lyase; *Gpat*, glycerol-3-phosphate O-acyltransferase; *Cpt2*, carnitine Palmitoyltransferase II; *Ldlr*, low density lipoprotein receptor. *P < 0.05; **P < 0.005.

**Suppl. Figure 8.**
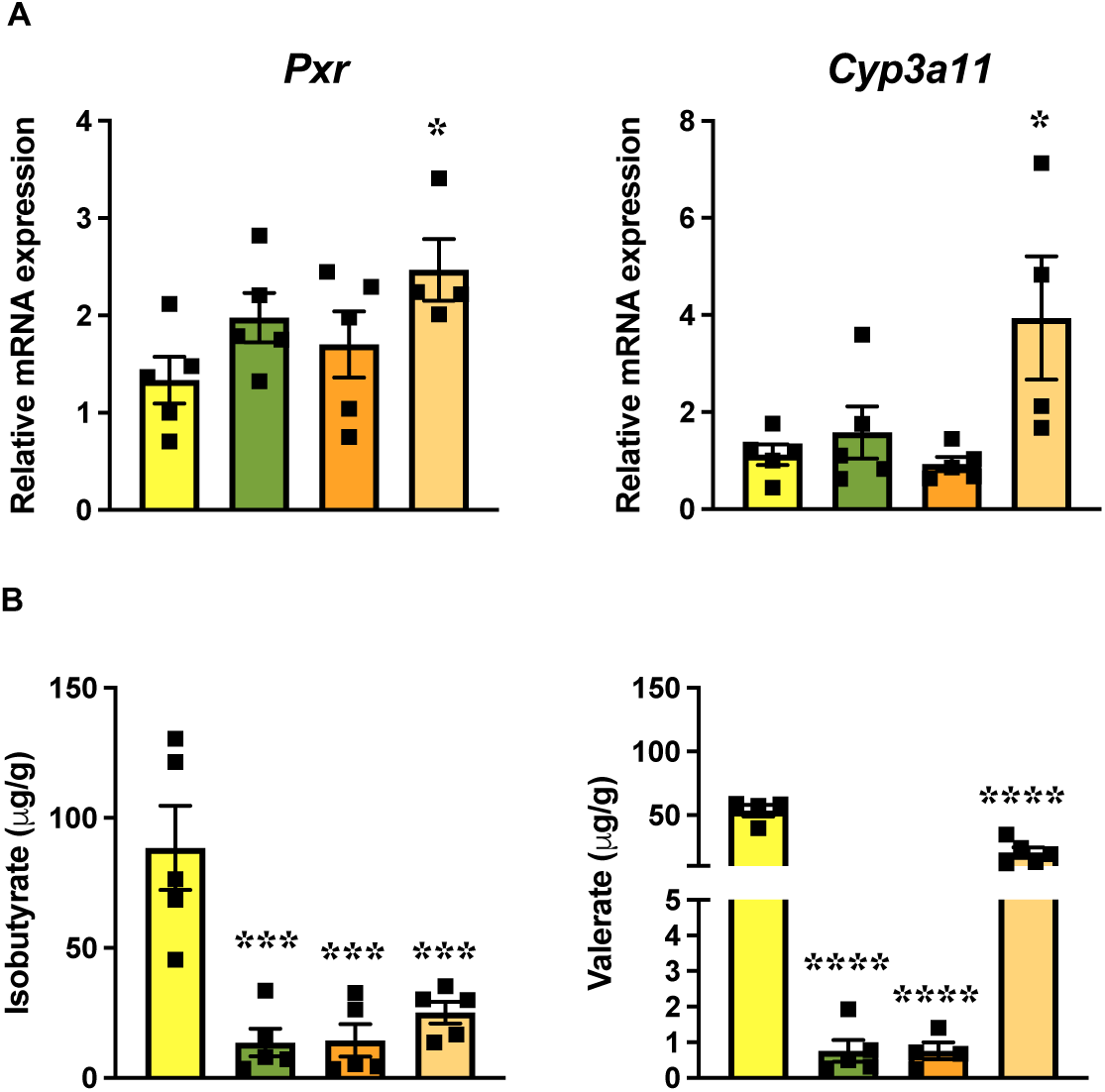
Differential effects of fibers on hepatic *Pxr* expression and gut microbial proteolytic fermentation. A) mRNA expression of Pxr and Cyp3a11 in livers, and B) levels of isobutyrate and valerate in cecal contents of WD and WD-FF fed mice. n=4-5 per group. Data are mean ± SEM; analyzed by 1-way analysis of variance with Sidak post hoc tests. *Cyp3a11*, cytochrome P 450 (CYP) family 3 subfamily A member 1; *Pxr*, pregnane X receptor. *P < 0.05; ***P < 0.001; ****P < 0.0001.

## References

1. Xie L, Alam MJ, Marques FZ, et al. A major mechanism for immunomodulation: Dietary fibres and acid metabolites. Seminars in Immunology 2023;66:101737.

2. Huang Z, Li Y, Park H, et al. Unveiling and harnessing the human gut microbiome in the rising burden of non-communicable diseases during urbanization. Gut Microbes 2023;15:2237645.

3. Ross FC, Patangia D, Grimaud G, et al. The interplay between diet and the gut microbiome: implications for health and disease. Nat Rev Microbiol 2024;22:671–686.

4. Levy M, Thaiss CA, Zeevi D, et al. Microbiota-Modulated Metabolites Shape the Intestinal Microenvironment by Regulating NLRP6 Inflammasome Signaling. Cell 2015;163:1428–43.

5. Dong TS, Guan M, Mayer EA, et al. Obesity is associated with a distinct brain-gut microbiome signature that connects Prevotella and Bacteroides to the brain’s reward center. Gut Microbes 2022;14:2051999.

6. Halfvarson J, Brislawn CJ, Lamendella R, et al. Dynamics of the human gut microbiome in inflammatory bowel disease. Nat Microbiol 2017;2:17004.

7. de Vos WM, Tilg H, Van Hul M, et al. Gut microbiome and health: mechanistic insights. Gut 2022;71:1020–1032.

8. Shelton CD, Byndloss MX. Gut Epithelial Metabolism as a Key Driver of Intestinal Dysbiosis Associated with Noncommunicable Diseases. Infect Immun 2020;88.

9. Zhao L, Zhang F, Ding X, et al. Gut bacteria selectively promoted by dietary fibers alleviate type 2 diabetes. Science 2018;359:1151–1156.

10. Lobel L, Cao YG, Fenn K, et al. Diet posttranslationally modifies the mouse gut microbial proteome to modulate renal function. Science 2020;369:1518–1524.

11. Byndloss MX, Bäumler AJ. The germ-organ theory of non-communicable diseases. Nature Reviews Microbiology 2018;16:103–110.

12. Vrieze A, Van Nood E, Holleman F, et al. Transfer of Intestinal Microbiota From Lean Donors Increases Insulin Sensitivity in Individuals With Metabolic Syndrome. Gastroenterology 2012;143:913–916.e7.

13. Witjes JJ, Smits LP, Pekmez CT, et al. Donor Fecal Microbiota Transplantation Alters Gut Microbiota and Metabolites in Obese Individuals With Steatohepatitis. Hepatol Commun 2020;4:1578–1590.

14. Stephen AM, Champ MM, Cloran SJ, et al. Dietary fibre in Europe: current state of knowledge on definitions, sources, recommendations, intakes and relationships to health. Nutr Res Rev 2017;30:149–190.

15. O’Keefe SJ. The association between dietary fibre deficiency and high-income lifestyle-associated diseases: Burkitt’s hypothesis revisited. Lancet Gastroenterol Hepatol 2019;4:984–996.

16. Gill SK, Rossi M, Bajka B, et al. Dietary fibre in gastrointestinal health and disease. Nature Reviews Gastroenterology & Hepatology 2021;18:101–116.

17. Deehan EC, Mocanu V, Madsen KL. Effects of dietary fibre on metabolic health and obesity. Nature Reviews Gastroenterology & Hepatology 2024;21:301–318.

18. Makki K, Deehan EC, Walter J, et al. The Impact of Dietary Fiber on Gut Microbiota in Host Health and Disease. Cell Host Microbe 2018;23:705–715.

19. Blaak EE, Canfora EE, Theis S, et al. Short chain fatty acids in human gut and metabolic health. Benef Microbes 2020;11:411–455.

20. Schroeder BO, Birchenough GMH, Ståhlman M, et al. Bifidobacteria or Fiber Protects against Diet-Induced Microbiota-Mediated Colonic Mucus Deterioration. Cell Host Microbe 2018;23:27–40.e7.

21. Zou J, Chassaing B, Singh V, et al. Fiber-Mediated Nourishment of Gut Microbiota Protects against Diet-Induced Obesity by Restoring IL-22-Mediated Colonic Health. Cell Host Microbe 2018;23:41–53.e4.

22. Wastyk HC, Fragiadakis GK, Perelman D, et al. Gut-microbiota-targeted diets modulate human immune status. Cell 2021;184:4137–4153.e14.

23. Rescigno M. Intestinal microbiota and its effects on the immune system. Cell Microbiol 2014;16:1004–13.

24. Coulon DB, Page R, Raggio AM, et al. Novel Resistant Starch Type 4 Products of Different Starch Origins, Production Methods, and Amounts Are Not Equally Fermented when Fed to Sprague-Dawley Rats. Mol Nutr Food Res 2020;64:e1900901.

25. Priyadarshini M, Navarro G, Reiman DJ, et al. Gestational Insulin Resistance Is Mediated by the Gut Microbiome-Indoleamine 2,3-Dioxygenase Axis. Gastroenterology 2022;162:1675–1689 e11.

26. Mina AI, LeClair RA, LeClair KB, et al. CalR: A Web-Based Analysis Tool for Indirect Calorimetry Experiments. Cell Metabolism 2018;28:656–666.e1.

27. Khan MW, Priyadarshini M, Cordoba-Chacon J, et al. Hepatic hexokinase domain containing 1 (HKDC1) improves whole body glucose tolerance and insulin sensitivity in pregnant mice. Biochim Biophys Acta Mol Basis Dis 2019;1865:678–687.

28. Lednovich KR, Nnyamah C, Gough S, et al. Intestinal FFA3 mediates obesogenic effects in mice on a Western diet. Am J Physiol Endocrinol Metab 2022;323:E290–e306.

29. Armstrong G, Martino C, Morris J, et al. Swapping Metagenomics Preprocessing Pipeline Components Offers Speed and Sensitivity Increases. mSystems 2022;7:e01378–21.

30. Zhu Q, Huang S, Gonzalez A, et al. Phylogeny-Aware Analysis of Metagenome Community Ecology Based on Matched Reference Genomes while Bypassing Taxonomy. mSystems 2022;7:e0016722.

31. Zhu Q, Mai U, Pfeiffer W, et al. Phylogenomics of 10,575 genomes reveals evolutionary proximity between domains Bacteria and Archaea. Nat Commun 2019;10:5477.

32. Langmead B, Salzberg SL. Fast gapped-read alignment with Bowtie 2. Nature Methods 2012;9:357–359.

33. Team RC. R: a language and environment for statistical computing. . R Foundation for Statistical Computing, Vienna, Austria., 2023.

34. Brennan C, Shaffer Justin P, Belda-Ferre P, et al. Streamlined extraction of nucleic acids and metabolites from low- and high-biomass samples using isopropanol and matrix tubes. Microbiology Spectrum 2025;13:e01912–25.

35. Bray JR, Curtis JT. An Ordination of the Upland Forest Communities of Southern Wisconsin. Ecological Monographs 1957;27:325–349.

36. Lozupone C, Lladser ME, Knights D, et al. UniFrac: an effective distance metric for microbial community comparison. The ISME Journal 2011;5:169–172.

37. UniProt: the universal protein knowledgebase. Nucleic Acids Res 2017;45:D158–d169.

38. Kanehisa M, Furumichi M, Tanabe M, et al. KEGG: new perspectives on genomes, pathways, diseases and drugs. Nucleic Acids Res 2017;45:D353–d361.

39. Röst HL, Sachsenberg T, Aiche S, et al. OpenMS: a flexible open-source software platform for mass spectrometry data analysis. Nat Methods 2016;13:741–8.

40. Anderson MJ. Permutational Multivariate Analysis of Variance (PERMANOVA). Wiley StatsRef: Statistics Reference Online:1–15.

41. McCarthy DJ, Chen Y, Smyth GK. Differential expression analysis of multifactor RNA-Seq experiments with respect to biological variation. Nucleic Acids Res 2012;40:4288–97.

42. Robinson MD, McCarthy DJ, Smyth GK. edgeR: a Bioconductor package for differential expression analysis of digital gene expression data. Bioinformatics 2010;26:139–40.

43. Ritchie ME, Phipson B, Wu D, et al. limma powers differential expression analyses for RNA-sequencing and microarray studies. Nucleic Acids Res 2015;43:e47.

44. Benjamini Y, Hochberg Y. Controlling the False Discovery Rate: A Practical and Powerful Approach to Multiple Testing. Journal of the Royal Statistical Society: Series B (Methodological) 1995;57:289–300.

45. Magne F, Gotteland M, Gauthier L, et al. The Firmicutes/Bacteroidetes Ratio: A Relevant Marker of Gut Dysbiosis in Obese Patients? Nutrients 2020;12.

46. Berry D. The emerging view of Firmicutes as key fibre degraders in the human gut. Environmental Microbiology 2016;18:2081–2083.

47. Dey P. All That Glitters Ain’t Gold: The Myths and Scientific Realities About the Gut Microbiota. Nutrients. Volume 17, 2025:3121.

48. Bailén M, Bressa C, Martínez-López S, et al. Microbiota Features Associated With a High-Fat/Low-Fiber Diet in Healthy Adults. Front Nutr 2020;7:583608.

49. Paudel D, Nair DVT, Tian S, et al. Dietary fiber guar gum-induced shift in gut microbiota metabolism and intestinal immune activity enhances susceptibility to colonic inflammation. Gut Microbes 2024;16:2341457.

50. Murga-Garrido SM, Hong Q, Cross T-WL, et al. Gut microbiome variation modulates the effects of dietary fiber on host metabolism. Microbiome 2021;9:117.

51. Paone P, Suriano F, Jian C, et al. Prebiotic oligofructose protects against high-fat diet-induced obesity by changing the gut microbiota, intestinal mucus production, glycosylation and secretion. Gut Microbes 2022;14:2152307.

52. Luis AS, Briggs J, Zhang X, et al. Dietary pectic glycans are degraded by coordinated enzyme pathways in human colonic Bacteroides. Nat Microbiol 2018;3:210–219.

53. Newman TM, Shively CA, Register TC, et al. Diet, obesity, and the gut microbiome as determinants modulating metabolic outcomes in a non-human primate model. Microbiome 2021;9:100.

54. Selmin OI, Papoutsis AJ, Hazan S, et al. n-6 High Fat Diet Induces Gut Microbiome Dysbiosis and Colonic Inflammation. International Journal of Molecular Sciences. Volume 22, 2021:6919.

55. Tang W, Ni Z, Wei Y, et al. Extracellular vesicles of Bacteroides uniformis induce M1 macrophage polarization and aggravate gut inflammation during weaning. Mucosal Immunology 2024;17:793–809.

56. Perez-Burillo S, Rajakaruna S, Paliy O. Growth of Bifidobacterium species is inhibited by free fatty acids and bile salts but not by glycerides. AIMS Microbiol 2022;8:53–60.

57. Kurdi P, Kawanishi K, Mizutani K, et al. Mechanism of growth inhibition by free bile acids in lactobacilli and bifidobacteria. J Bacteriol 2006;188:1979–86.

58. Zünd JN, Mujezinovic D, Reichlin M, et al. Novel cross-feeding human gut microbes metabolizing tryptophan to indole-3-propionate. Gut Microbes 2025;17:2501195.

59. Zhang SM, Wu HC, Hung JH, et al. Indole Derivatives Biosynthesis in Bifidobacterium longum subsp. infantis and the Tryptophan Substrate Availability. Microb Biotechnol 2025;18:e70167.

60. Yong CC, Sakurai T, Kaneko H, et al. Human gut-associated Bifidobacterium species salvage exogenous indole, a uremic toxin precursor, to synthesize indole-3-lactic acid via tryptophan. Gut Microbes 2024;16:2347728.

61. Sinha AK, Laursen MF, Brinck JE, et al. Dietary fibre directs microbial tryptophan metabolism via metabolic interactions in the gut microbiota. Nature Microbiology 2024;9:1964–1978.

62. Devlin AS, Marcobal A, Dodd D, et al. Modulation of a Circulating Uremic Solute via Rational Genetic Manipulation of the Gut Microbiota. Cell Host Microbe 2016;20:709–715.

63. Laurans L, Venteclef N, Haddad Y, et al. Genetic deficiency of indoleamine 2,3-dioxygenase promotes gut microbiota-mediated metabolic health. Nat Med 2018;24:1113–1120.

64. Natividad JM, Agus A, Planchais J, et al. Impaired Aryl Hydrocarbon Receptor Ligand Production by the Gut Microbiota Is a Key Factor in Metabolic Syndrome. Cell Metabolism 2018;28:737–749.e4.

65. Kwon YH, Khan WI. Peripheral serotonin: cultivating companionship with gut microbiota in intestinal homeostasis. Am J Physiol Cell Physiol 2022;323:C550–c555.

66. Agus A, Clément K, Sokol H. Gut microbiota-derived metabolites as central regulators in metabolic disorders. Gut 2021;70:1174–1182.

67. Roager HM, Licht TR. Microbial tryptophan catabolites in health and disease. Nat Commun 2018;9:3294.

68. Tennoune N, Andriamihaja M, Blachier F. Production of Indole and Indole-Related Compounds by the Intestinal Microbiota and Consequences for the Host: The Good, the Bad, and the Ugly. Microorganisms 2022;10.

69. Forsyth CB, Voigt RM, Keshavarzian A. Intestinal CYP2E1: A mediator of alcohol-induced gut leakiness. Redox Biol 2014;3:40–6.

70. Armand L, Fofana M, Couturier-Becavin K, et al. Dual effects of the tryptophan-derived bacterial metabolite indole on colonic epithelial cell metabolism and physiology: comparison with its co-metabolite indoxyl sulfate. Amino Acids 2022;54:1371–1382.

71. Cho YE, Kim DK, Seo W, et al. Fructose Promotes Leaky Gut, Endotoxemia, and Liver Fibrosis Through Ethanol-Inducible Cytochrome P450-2E1-Mediated Oxidative and Nitrative Stress. Hepatology 2021;73:2180–2195.

72. Martinez-Medina M, Denizot J, Dreux N, et al. Western diet induces dysbiosis with increased E coli in CEABAC10 mice, alters host barrier function favouring AIEC colonisation. Gut 2014;63:116–24.

73. Shashikanth N, France MM, Xiao R, et al. Tight junction channel regulation by interclaudin interference. Nat Commun 2022;13:3780.

74. Pircalabioru G, Aviello G, Kubica M, et al. Defensive Mutualism Rescues NADPH Oxidase Inactivation in Gut Infection. Cell Host & Microbe 2016;19:651–663.

75. Qiu X, Macchietto MG, Liu X, et al. Identification of gut microbiota and microbial metabolites regulated by an antimicrobial peptide lipocalin 2 in high fat diet-induced obesity. Int J Obes (Lond) 2021;45:143–154.

76. Chen X, Fu L, Zhu Z, et al. Exploring the association between dietary fiber intake and hepatic steatosis: insights from NHANES. BMC Gastroenterology 2024;24:160.

77. Régnier M, Carbinatti T, Parlati L, et al. The role of ChREBP in carbohydrate sensing and NAFLD development. Nat Rev Endocrinol 2023;19:336–349.

78. Bakr AF, Farag MA. Soluble Dietary Fibers as Antihyperlipidemic Agents: A Comprehensive Review to Maximize Their Health Benefits. ACS Omega 2023;8:24680–24694.

79. Skytthe M, Graversen JH, Moestrup SK. CD163 Expression Protects Against Early Hepatic Steatosis in Western Diet-Fed Male Mice. Gastro Hep Advances 2026;5.

80. Kang J, Postigo-Fernandez J, Kim K, et al. Notch-mediated hepatocyte MCP-1 secretion causes liver fibrosis. JCI Insight 2023;8.

81. Banoglu E, King RS. Sulfation of indoxyl by human and rat aryl (phenol) sulfotransferases to form indoxyl sulfate. Eur J Drug Metab Pharmacokinet 2002;27:135–40.

82. Gong P, Cederbaum AI. Transcription Factor Nrf2 Protects HepG2 Cells against CYP2E1 plus Arachidonic Acid-dependent Toxicity*. Journal of Biological Chemistry 2006;281:14573–14579.

83. Zhou J, Zheng Q, Chen Z. The Nrf2 Pathway in Liver Diseases. Frontiers in Cell and Developmental Biology 2022;Volume 10 - 2022.

84. Togao M, Asakawa N, Wagai G, et al. Dual effects of indoxyl sulfate on modulation of human hepatic CYP3A activity, with individual differences. PLOS ONE 2025;20:e0328182.

85. Sári Z, Mikó E, Kovács T, et al. Indoxylsulfate, a Metabolite of the Microbiome, Has Cytostatic Effects in Breast Cancer via Activation of AHR and PXR Receptors and Induction of Oxidative Stress. Cancers. Volume 12, 2020:2915.

86. Müller M, Hernández MAG, Goossens GH, et al. Circulating but not faecal short-chain fatty acids are related to insulin sensitivity, lipolysis and GLP-1 concentrations in humans. Sci Rep 2019;9:12515.

87. Brockman DA, Chen X, Gallaher DD. High-Viscosity Dietary Fibers Reduce Adiposity and Decrease Hepatic Steatosis in Rats Fed a High-Fat Diet. The Journal of Nutrition 2014;144:1415–1422.

88. Armstrong HK, Bording-Jorgensen M, Santer DM, et al. Unfermented β-fructan Fibers Fuel Inflammation in Select Inflammatory Bowel Disease Patients. Gastroenterology 2023;164:228–240.

89. Priyadarshini M, Kotlo KU, Dudeja PK, et al. Role of Short Chain Fatty Acid Receptors in Intestinal Physiology and Pathophysiology. Compr Physiol 2018;8:1091–1115.

90. Dugas LR, Bernabé BP, Priyadarshini M, et al. Decreased microbial co-occurrence network stability and SCFA receptor level correlates with obesity in African-origin women. Sci Rep 2018;8:17135.

91. Calleja-Conde J, Bühler K-M, Echeverry-Alzate V, et al. Fermentable dietary fibers reduce voluntary alcohol intake and modulate gut microbiota composition in rats. Journal of Functional Foods 2025;134:107079.

92. Kennedy MS, Freiburger A, Cooper M, et al. Diet outperforms microbial transplant to drive microbiome recovery in mice. Nature 2025;642:747–755.

93. Holscher HD. Dietary fiber and prebiotics and the gastrointestinal microbiota. Gut Microbes 2017;8:172–184.

94. Miles JP, Zou J, Kumar M-V, et al. Supplementation of Low- and High-fat Diets with Fermentable Fiber Exacerbates Severity of DSS-induced Acute Colitis. Inflammatory Bowel Diseases 2017;23:1133–1143.

95. Qi Q, Li J, Yu B, et al. Host and gut microbial tryptophan metabolism and type 2 diabetes: an integrative analysis of host genetics, diet, gut microbiome and circulating metabolites in cohort studies. Gut 2022;71:1095.

96. Tuomainen M, Lindström J, Lehtonen M, et al. Associations of serum indolepropionic acid, a gut microbiota metabolite, with type 2 diabetes and low-grade inflammation in high-risk individuals. Nutrition & Diabetes 2018;8:35.

97. de Mello VD, Paananen J, Lindström J, et al. Indolepropionic acid and novel lipid metabolites are associated with a lower risk of type 2 diabetes in the Finnish Diabetes Prevention Study. Sci Rep 2017;7:46337.

98. Laursen MF, Sakanaka M, von Burg N, et al. Bifidobacterium species associated with breastfeeding produce aromatic lactic acids in the infant gut. Nat Microbiol 2021;6:1367–1382.

99. Cervantes-Barragan L, Chai JN, Tianero MD, et al. Lactobacillus reuteri induces gut intraepithelial CD4(+)CD8αα(+) T cells. Science 2017;357:806–810.

100. Aragozzini F, Ferrari A, Pacini N, et al. Indole-3-lactic acid as a tryptophan metabolite produced by Bifidobacterium spp. Appl Environ Microbiol 1979;38:544–6.

101. Choi BSY, Daniel N, Houde VP, et al. Feeding diversified protein sources exacerbates hepatic insulin resistance via increased gut microbial branched-chain fatty acids and mTORC1 signaling in obese mice. Nature Communications 2021;12:3377.

102. Shaheen N, Miao J, Xia B, et al. Multifaceted Role of Microbiota-Derived Indole-3-Acetic Acid in Human Diseases and Its Potential Clinical Application. Faseb j 2025;39:e70574.

103. Ding Y, Yanagi K, Yang F, et al. Oral supplementation of gut microbial metabolite indole-3-acetate alleviates diet-induced steatosis and inflammation in mice: eLife Sciences Publications, Ltd, 2023.

104. Lin T-Y, Wu W-K, Hung S-C. High interindividual variability of indoxyl sulfate production identified by an oral tryptophan challenge test. npj Biofilms and Microbiomes 2025;11:15.

