## Supplementary table 1 for "A high fermentable fiber Western diet reduces indole levels"

**Supplemental Table 1** **Composition of control, western and fiber supplemented custom formulated diets**

| Product # | D19010908 | | D19070209 | | D19070210 | | D19070211 | | D19070212 | |
|---|---|---|---|---|---|---|---|---|---|---|
| % | gm | *kcal* | gm | *kcal* | gm | *kcal* | gm | *kcal* | gm | *kcal* |
| Protein | 17 | ***17*** | 20 | ***17*** | 17 | ***17*** | 17 | ***17*** | 18 | ***17*** |
| Carbohydrate | 71 | ***73*** | 50 | ***43*** | 56 | ***43*** | 56 | ***43*** | 55 | ***43*** |
| Fat | 4 | ***10*** | 21 | ***40*** | 18 | ***40*** | 18 | ***40*** | 19 | ***40*** |
| Total | | *100* | | *100* | | *100* | | *100* | | *100* |
| kcal/gm | 3.9 | | 4.7 | | 4.1 | | 4.1 | | 4.2 | |
| Ingredient | gm | *kcal* | gm | *kcal* | gm | *kcal* | gm | *kcal* | gm | *kcal* |
| Casein | 195 | *780* | 195 | *780* | 195 | *780* | 195 | *780* | 195 | *780* |
| DL-Methionine | 3 | *12* | 3 | *12* | 3 | *12* | 3 | *12* | 3 | *12* |
| Corn Starch | 0 | *0* | 0 | *0* | 0 | *0* | 0 | *0* | 0 | *0* |
| Amioca | 695 | *2780* | 109.5 | *438* | 23.25 | *93* | 18 | *72* | 0 | *0* |
| Maltodextrin 10 | 150 | *600* | 100 | *400* | 100 | *400* | 100 | *400* | 100 | *400* |
| Sucrose | 0 | *0* | 281.5 | *1126* | 281.5 | *1126* | 281.5 | *1126* | 281.5 | *1126* |
| Cellulose, BW200 | 50 | *0* | 50 | *0* | 50 | *0* | 50 | *0* | 50 | *0* |
| Fructo-oligosaccharides | 0 | *0* | 0 | *0* | 230 | *345* | 0 | *0* | 0 | *0* |
| Pectin | 0 | *0* | 0 | *0* | 0 | *0* | 228 | *364.8* | 0 | *0* |
| Guar Gum | 0 | *0* | 0 | *0* | 0 | *0* | 0 | *0* | 225 | *438.75* |
| Milk Fat, Anhydrous | 42.5 | *382.5* | 200 | *1800* | 200 | *1800* | 200 | *1800* | 200 | *1800* |
| Corn Oil | 10 | *90* | 10 | *90* | 10 | *90* | 10 | *90* | 10 | *90* |
| Ethoxyquin | 0.04 | *0* | 0.04 | *0* | 0.04 | *0* | 0.04 | *0* | 0.04 | *0* |
| Mineral Mix S10001 | 35 | *0* | 35 | *0* | 35 | *0* | 35 | *0* | 35 | *0* |
| Calcium Carbonate | 4 | *0* | 4 | *0* | 4 | *0* | 4 | *0* | 4 | *0* |
| Vitamin Mix V10001 | 10 | *40* | 10 | *40* | 10 | *40* | 10 | *40* | 10 | *40* |
| Choline Bitartrate | 2 | *0* | 2 | *0* | 2 | *0* | 2 | *0* | 2 | *0* |
| Cholesterol | 0 | *0* | 1.5 | *0* | 1.5 | *0* | 1.5 | *0* | 1.5 | *0* |
| FD&C Yellow Dye #5 | 0 | *0* | 0 | *0* | 0.05 | *0* | 0 | *0* | 0.04 | *0* |
| FD&C Red Dye #40 | 0.05 | *0* | 0 | *0* | 0 | *0* | 0 | *0* | 0 | *0* |
| FD&C Blue Dye #41 | 0 | *0* | 0 | *0* | 0 | *0* | 0.05 | *0* | 0.01 | *0* |
| **Total** | **1196.59** | ***4685*** | **1001.54** | ***4686*** | **1145.34** | ***4686*** | **1138.09** | ***4685*** | **1117.09** | ***4687*** |
| **% Fermentable Fiber** | | | | | | | | | | |
| Fructo-oligosaccharides | 0.0 | | 0.0 | | 20.1 | | 0.0 | | 0.0 | |
| Pectin | 0.0 | | 0.0 | | 0.0 | | 20.0 | | 0.0 | |
| Guar Gum | 0.0 | | 0.0 | | 0.0 | | 0.0 | | 20.1 | |
