## Supplementary table 2 for "A high fermentable fiber Western diet reduces indole levels"

**Supplemental Table 2** **Primer sequences used for quantitative polymerase chain reaction**

| Gene | Forward primer 5'-3' | Reverse primer 5'-3' |
|---|---|---|
| *Il-6* | TAGTCCTTCCTACCCCAATTTCC | TTGGTCCTTAGCCACTCCTTC |
| *Ccl2* | TTAGCCCTGACCGAGAAAGA | AAAGGACCTCTCTGGTGCTG |
| *Ccl-3* | TTCTCTGTACCATGACACTCTGC | CGTGGAATCTTCCGGCTGTAG |
| *Tnfa* | GGTGCCTATGTCTCAGCCTCTT | GCCATAGAACTGATGAGAGGGAG |
| *Lcn2* | CCTCCATCCTGGTCAGGGAC | TAGTCCGTGGTGGCCACTTG |
| *F4/80* | GATGTGGAGGATGGGAGATG | ACAGCAGGAAGGTGGCTATG |
| CD163 | GGCTAGACGAAGTCATCTGCAC | CTTCGTTGGTCAGCCTCAGAGA |
| Vsig4 | ATGTGAGGTCACCTGGCAGACT | GCAGGGTTGTAGGTGCTTCAGT |
| CD68 | GGCGGTGGAATACAATGTGTCC | AGCAGGTCAAGGTGAACAGCTG |
| *Claudin-4* | AGCAAACGTCCACTGTCCTT | AATCCACCTCCACCCTTCTT |
| *Claudin-2* | TTAGCCCTGACCGAGAAAGA | AAAGGACCTCTCTGGTGCTG |
| *Ahr* | CTTTGCTGAACTCGGCTTGC | TTGCTGGGGGCACACCATCT |
| *Pxr* | AGAGATCATCCCTCTTCTGCCAC | GATCTGGTCCTCAATAGGCAGGT |
| *Cyp1a1* | GGGTTTGACACAGTCACAA | GGGACGAAGGATGAATGCC |
| *Cyp3a11* | TCACACACACAGTTGTAGGCAGAA | GTTTACGAGTCCCATATCGGTAGAG |
| *Cyp2e1* | CGTTGCCTTGCTTGTCTGGA | AAGAAAGGAATTGGGAAAGGTCC |
| *Sult1a1* | CACAAGGGTCCTCTCCTTAGC | TGACAGCGGAACGTGAAGTC |
| *Nrf2* | TCTTGGAGTAAGTCGAGAAGTGT | GTTGAAACTGAGCGAAAAAGGC |
| *Ho-1* | AAGCCGAGAATGCTGAGTTCA | GCCGTGTAGATATGGTACAAGGA |
| *Nqo1* | ATGGGAGGTGGTCGAATCTGA | GCCTTCCTTATACGCCAGAGATG |
| *Nox-1* | GGTTGGGGCTGAACATTTTTC | TCGACACACAGGAATCAGGAT |
| *Nox-4* | GAAGGGGTTAAACACCTCTGC | ATGCTCTGCTTAAACACAATCCT |
| *Cox2* | TTCAACACACTCTATCACTGGC | AGAAGCGTTTGCGGTACTCAT |
| *Acly* | CCTTGAAGGAAGCAGGAGTG | GGTCATGAATGAGGCAGGTT |
| *Srebp1c* | GGAGCCATGGATTGCACATT | GGCCCGGGAAGTCACTGT |
| *Fasn* | TGAGCACACTGCTGGTGAAC | CAGGTTCGGAATGCTATCCA |
| *Gpat* | ACAGTTGGCACAATAGACGTTT | CCTTCCATTTCAGTGTTGCAGA |
| *Chrebp1a* | CGACACTCACCCACCTCTTC | TTGTTCAGCCGGATCTTGTTC |
| *Chrebp1b* | TCTGCAGATCGCGTGGAG | CTTGTCCCGGCATAGCAAC |
| *Ldlr* | TGTCACCTGTCAGTCCAATCAA | TCAGAGCCATCTAGGCAATCTC |
| *Acc1* | ATCCTGCGAACCTGGATTCT | CCCACCAGAGAAACCTCTCC |
| *Cd36* | GGACATACTTAGATGTGGAACCCATA | TGTTGACCTGCAGTCGTTTTG |
| *Cpt2* | GGATAAACAGAATAAGCACACCA | GAAGGAACAAAGCGGATGAG |
| *B actin* | TAC CAC AGG CAT TGT GAT GG | TCT CAG CTG TGG TGG TGA AG |
| *Gapdh* | AGCTTGTCATCAACGGGAAG | TTTGATGTTAGTGGGGTCTCG |
